# Dynamic Profiling of Binding and Allosteric Propensities of the SARS-CoV-2 Spike Protein with Different Classes of Antibodies: Mutational and Perturbation-Based Scanning Reveal Allosteric Duality of Functionally Adaptable Hotspots

**DOI:** 10.1101/2021.04.13.439743

**Authors:** Gennady M. Verkhivker, Steve Agajanian, Deniz Yazar Oztas, Grace Gupta

## Abstract

Structural and biochemical studies of the SARS-CoV-2 spike complexes with highly potent antibodies have revealed multiple conformation-dependent epitopes and a broad range of recognition modes linked to different neutralization responses In this study, we combined atomistic simulations with mutational and perturbation-based scanning approaches to perform in silico profiling of binding and allosteric propensities of the SARS-CoV-2 spike protein residues in complexes with B38, P2B-2F6, EY6A and S304 antibodies representing three different classes. Conformational dynamics analysis revealed that binding-induced modulation of soft modes can elicit the unique protein response to different classes of antibodies. Mutational scanning heatmaps and sensitivity analysis revealed the binding energy hotspots for different classes of antibodies that are consistent with the experimental deep mutagenesis, showing that differences in the binding affinity caused by global circulating variants in spike positions K417, E484 and N501 are relatively moderate and may not fully account for the observed antibody resistance effects. Through functional dynamics analysis and perturbation-response scanning of the SARS-CoV-2 spike protein residues in the unbound form and antibody-bound forms, we examine how antibody binding can modulate allosteric propensities of spike protein residues and determine allosteric hotspots that control signal transmission and global conformational changes. These results show that residues K417, E484, and N501 targeted by circulating mutations correspond to a group of versatile allosteric centers in which small perturbations can modulate collective motions, alter the global allosteric response and elicit binding resistance. We suggest that SARS-CoV-2 S protein may exploit plasticity of specific allosteric hotspots to generate escape mutants that alter response to antibody binding without compromising activity of the spike protein.

## Introduction

The structural and biochemical studies of the SARS-CoV-2 spike (S) glycoproteins and complexes with highly potent antibodies have revealed multiple conformation-dependent epitopes highlighting the link between conformational plasticity of spike proteins and capacity for eliciting specific binding and broad neutralization responses.^1–7^ The potent antibodies often compete and interfere with the mechanism of virus infection that features spontaneous conformational transformations of the SARS-CoV-2 S protein between closed “down” and open “up” forms of the receptor-binding domain (RBD).^7–9^ A general mechanism of population shifts between different functional states of the SARS-CoV-2 S trimers suggested that RBD epitopes can become stochastically exposed to the interactions with the host receptor ACE, leading to the increased population of open S-RBD states priming spike protein for fusion activation.^10–16^ The initial structure of SARS-CoV-RBD complex with a neutralizing antibody 80R showed that the epitope on the S1 RBD overlapped closely with the ACE2-binding site, suggesting that a direct interference mechanism may be responsible for the neutralizing activity.^17^ However, several SARS-CoV–specific neutralizing antibodies such as m396, 80R, and F26G19 that block the RBM motif in the open S conformation did not exhibit a strong neutralizing activity against SARS-CoV-2 protein. A wide spectrum of SARS-CoV-2 antibodies can be divided into several main classes of which class I and class II antibodies target epitopes that target the receptor binding motif (RBM) region of the RBD.^4–6^ Class I is the largest group of structurally characterized monoclonal antibodies against the SARSCoV-2 S-RBD that exhibit a significant overlap with the ACE2 epitope and bind when the RBD is in the open “up” state. Structural and biochemical studies characterized the binding epitopes and molecular mechanisms for a number of class I SARS-CoV-2 antibodies including REGN10933,^18^ B38,^19^ CB6,^20^ CC12.3,^21^ C105,^7^ and BD-236.^22^ The REGN-COV2 cocktail of REG10933 (RBM-specific class I) and REGN10987 (RBD core-specific) demonstrated significant potential in preventing mutational escape.^23^ SARS-CoV-2 antibodies B38 and H4 can bind simultaneously to different epitopes on RBD such that both antibodies together confer a stronger neutralizing effect than either antibody individually. Structural analysis showed that B38 binding epitope spans 36 residues on the RBD of which 15 are conserved between SARS-Cov-2 and SARS-CoV and reside within the ACE2 binding interface.^19^ Some of the known escaping mutations are within the epitopes for the neutralizing antibody B38 (K458R, D405V).^24^ A new SARS-CoV-2 lineage (501Y.V2) first detected in South Africa is characterized by 21 mutations with 8 lineage-defining mutations in the S protein, including three at important RBD residues (K417N, E484K and N501Y) that have functional significance and often induce significant immune escape.^25, 26^ The latest analysis of 17 class I antibody structures confirmed conservation of their conformational epitopes centered on RBD residue K417 which is one of three substitutions in the RBD of the circulating mutation of 501Y.V2 lineage.^27^ The antibodies potently neutralized the original lineage, but not the 501Y.V2 pseudovirus revealing a strong dependence on the K417 residue. Class II neutralizing antibodies P2B-2F6^28^, BD-368-2^22^, CV07-270,^29^ S2H13^30^, and C121^6^ revealed the epitopes that only partially overlap with the ACE2 binding site and, as a result, these antibodies can bind to the S-RBD in both up and down conformation. The potential neutralization mechanisms for class II are more diverse and can include blocking ACE2 from binding and also inducing S1 domain shedding by trapping the RBDs in the up state. Structural comparison of 15 class II antibodies revealed key interactions with residue E484, showing most of these antibodies can be evaded by 501Y.V2 RBD variant that prominently featured E484K mutation.^27^ Crystal structure of the S-RBD complex with P2B-2F6 revealed only moderate steric hindrance with the ACE2 binding site and partial interference with viral engagement with ACE2 recepotor.^28^ Of 12 RBM contacts, only four residues (N448, G447, Y449, and N450) are conserved in SARS-CoV-2 and SARS-CoV proteins while the only common residues recognized by both P2B-2F6 and ACE2 are G446 and Y449. Despite a small number of key overlapping residues, similar high binding affinities of P2B-2F6 and ACE2 binding suggest that they compete for the RBD interactions.^28^ Recently, the crystal structures of several other related antibodies P2C-1F11 and P2C-1A3 bound to the SARS-CoV-2 RBD were determined, showing instructive differences in the angles of antibody approach the RBD and highlighting common and unique binding residues on the S-RBD.^31^ Of special interest, one antibody, P2C-1F11 that most closely mimics binding of receptor ACE2, displays the most potent neutralizing activity in vitro and demonstrates the highest binding affinity to RBD. Functional studies showed that F490L, V483A an L452R mutations can reduce considerably sensitivity to P2B2F6 antibody.^24^ Analysis of the molecular determinants and mechanisms of mutational escape showed that SARS-CoV-2 virus rapidly escapes from individual antibodies but doesn’t easily escape from the cocktail due to stronger evolutionary constraints on RBD-ACE2 interaction and RBD protein folding.^32^

Class III antibodies including CR3022^33–35^ EY6A^36^, and S304^30^ bind to the opposite face of the RBD targeting the cryptic epitope that is only accessible when at least two RBDs are in the up state. Structural and surface plasmon resonance studies confirmed that CR3022 binds the RBD of SARS-CoV-2 displaying strong neutralization by allosterically perturbing the interactions between the RBD regions and ACE2 receptor.^34^ The neutralization mechanism of SARS-CoV-2 through destabilization of the prefusion S conformation can provide a resistance mechanism to virus escape which can be contrasted with class I antibodies directly competing for the ACE2-binding site and often susceptible to immune evasion. EY6A antibody binds to the same cryptic epitope as CR3022 but exploit a different orientation with respect to the S-RBD corresponding to a 73° rotation around an axis perpendicular to the RBD α3-helix.^36^ Interestingly, EY6A binding to the isolated RBD is an order of magnitude better than CR3022 and the unique binding mode allows three EY6A molecules to bind simultaneously around the central axis of the S protein.^36^ The antibodies isolated from 10 convalescent COVID-19 patients showed neutralizing activities against authentic SARS-CoV-2, with 4A8 antibody binding to the NTD of the S protein conformation with one RBD in “up” conformation and the other two RBDs in “down” conformation.^37^ Potent neutralizing antibodies from COVID-19 patients examined through electron microscopy studies confirmed that the SARS-CoV-2 S protein features multiple distinct antigenic sites, including RBD-based and non-RBD epitopes.^38^ Cryo–EM characterization of the SARS-CoV-2 S trimer in complex with the H014 Fab fragment revealed a new conformational epitope that is accessible only when the RBD is in the up conformation.^39^ Biochemical and virological studies demonstrated that H014 prevents attachment of SARS-CoV-2 to the host cell receptors and can exhibit broad cross-neutralization activities by leveraging conserved nature of the RBD epitope and a partial overlap with ACE2-binding region. The recently reported S309 antibody potently neutralizes both SARS-CoV-2 and SARS-CoV through binding to a conserved RBD epitope which is distinct from the RBM region and accessible in both open and closed states, so that there is no completion between S309 and ACE2 for binding to the SARS-CoV-2 S protein.^40^ Two ultra-potent Abs S2M11 and S2E12 targeting the overlapping RBD epitopes were recently reported, revealing antibody-specific modulation of protein responses and adaptation of different functional states for the S trimer.^41^ Cryo-EM structures showed that S2M11 can recognize and stabilize S protein in the closed conformation by binding to a quaternary epitope spanning two RBDs of the adjacent protomers in the S trimer, while S2E12 binds to a tertiary epitope contained within one S protomer and shifts the conformational equilibrium towards a fully open S trimer conformation. The emerging body of studies suggested that properly designed antibody cocktails can provide a broad and efficient cross-neutralization effects through synergistic targeting of conserved and more variable SARS-CoV-2 RBD epitopes, thereby offering a robust strategy to combat virus resistance. These studies also suggested that several potent antibodies may function by allosterically interfering with the host receptor binding and causing conformational changes in the S protein that can obstruct other epitopes and block virus infection without directly interfering with ACE2 recognition.

Biochemical and functional studies using a protein engineering and deep mutagenesis have quantified binding mechanisms of SARS-CoV-2 interactions with the host receptor.^42, 43^ Deep mutational scanning of SARS-CoV-2 RBD revealed mechanisms of SARS-CoV-2 interactions with the host receptor showing that many mutations of the RBD residues can be well tolerated with respect to both folding and binding. A number of amino acid modifications could even improve ACE2 binding, including important binding interface positions that enhance RBD expression (V367F and G502D) or enhance ACE2 affinity (N501F, N501T, and Q498Y).^42^ Functional studies characterized the key amino acid residues of the RBD for binding with human ACE2 and neutralizing antibodies, revealing two groups of amino acid residues that modulate binding, where the SARS-CoV-2 RBD mutations in positions N439, L452, T470, E484, Q498 and N501 can result in the enhanced binding affinity for ACE2.^44^ Interestingly, residues E484 and F486 in the SARS-CoV-2 RBD were identified as important sites for the recognition and differential binding ACE2 and neutralizing antibodies, indicating that structural plasticity of the RBM residues may induce mutational escape from the RBD-targeting neutralizing antibodies. Functional mapping of mutations in the SARS-CoV-2 S-RBD using deep mutational scanning showed that escape sites from antibodies can be constrained to ensure proper expression of the folded RBD, suggesting that escape-resistant antibody cocktails can compete for binding to the same RBD region but have different escape mutations.^45^ Comprehensive mapping of mutations in the SARS-CoV-2 RBD and neutralization assay experiments indicated that E484 modifications can reduce the neutralization potency by some antibodies by >10 fold.^47–51^ REGN10933 and REGN10987 are escaped by different mutations as mutation at F486 escaped neutralization only by REGN10933, whereas mutations at K444 escaped neutralization only by REGN10987, while only a single mutation E406W escaped the REGN-COV2 cocktail (REG10987+REG10933).^49, 50^ These studies indicated that mutations in the epitope centered around E484 position (G485, F486, F490) or in the 443–455 loop (K444, V445, L455, F456 sites) strongly affected neutralization for different classes of antibodies. By combining biophysical experiments with crystallographic studies, the binding epitopes for 80 monoclonal antibodies that bind the RBD was determined.^52^ The resulting map shows the antibody footprints forming five major epitopes by cluster analysis, particularly highlighting the role residues E484–F486 that bridge the epitopes and are accessible to antibodies from different approaching angles of attack. Notably, mutations in these positions E484K and F486L were identified as recurrent mutations identified in the B.1.351 and P.1 lineages.^53, 54^

Computational modeling and molecular dynamics (MD) simulations have examined conformational and energetic mechanisms of SARS-CoV-2 functions.^55–60^ All-atom MD simulations of the SARS-CoV-2 S protein confirmed dynamic fluctuations between open and closed spike states by constructing the free energy landscapes and minimum energy pathways, also revealing that RBD switches to the up position through an obligatory semi-open intermediate that reduces the free energy barrier between functional forms and could serve as a prerequisite state for the host cell recognition.^55^ MD simulations of the full-length SARS-CoV-2 S glycoprotein embedded in the viral membrane, with a complete glycosylation profile were recently reported, providing the unprecedented level of details about open and closed structures.^57^ A comprehensive study employed MD simulations to reveal a balance of hydrophobic interactions and elaborate hydrogen-bonding network in the SARS-CoV-2-RBD interface.^59^ Computational studies of the SARS-CoV-2 S trimer interactions with ACE2 using the recent crystal structures^61–64^ also provided important insights into the key determinants of the binding affinity and selectivity. Molecular mechanisms of the SARS-CoV-2 binding with ACE2 were analyzed in our recent study using coevolution and conformational dynamics.^63^ A series of all-atom MD simulations of the SARS-CoV-2 S-RBDS complex with ACE2 in the absence and presence of external force examined the effects of alanine substitutions showing that the hydrophobic end of RBD serves as the main energetic hotspot for ACE2 binding.^64^ By using molecular simulations and network modeling we recently presented evidence that the SARS-CoV-2 spike protein can function as an allosteric regulatory engine that fluctuates between dynamically distinct functional states.^65^ Coarse-grained normal mode analyses combined with Markov model and computation of transition probabilities characterized the dynamics of the S protein and the effects of mutational variants D614G and N501Y on protein dynamics and energetics.^66^ Using time-independent component analysis and protein networks, another computational study identified the hotspot residues that may exhibit long-distance coupling with the RBD opening, showing that some mutations may allosterically affect the stability of the RBD regions.^67^ Molecular simulations reveal that N501Y mutation increases ACE2 binding affinity, and may impact the collective dynamics of the ACE2-RBD complex while mutations K417N and E484K reduce the ACE2-binding affinity.^68^

In this study, we combined MD simulations with mutational and perturbation-based scanning approaches to perform a comprehensive structural profiling of binding and allosteric propensities of the SARS-CoV-2 S-RBD residues in complexes with three different classes of antibodies. We investigated dynamics, energetics and allosteric potential of the S-RBD protein in complexes with B38, P2B-2F6, EY6A and S304 antibodies. The mutational scanning-based heatmaps reveal the binding energy hotspots for different classes of antibodies indicating that differences in the binding affinity caused by global variants are moderate and are unlikely to fully account for the antibody resistance effects. Through functional dynamics analysis and perturbation-response scanning of the SARS-CoV-2 S-RBD in the unbound form and antibody-bound forms, we characterize allosteric propensities of spike protein residues and determine regulatory sites that may function as allosteric hotspots of long-range communications. These results show that residues K417, E484, and N501 targeted by circulating mutations correspond to a group of allosteric centers involved in a coordinated cross-talk to enable antibody-induced modulation of the spike activity. Using different classes of antibodies, we show that mutational variants constrained by the requirements for preservation of the RBD stability may preferentially target a group of structurally adaptable allosteric centers in which small perturbations can modulate functional motions, alter the global allosteric response and elicit binding resistance. We suggest that SARS-CoV-2 S protein may exploit plasticity of specific allosteric hotspots to generate escape mutants that alter response to antibody binding without compromising activity of the spike protein.

## Materials and Methods

### Structure Preparation and Analysis

The structures of the SARS-CoV-RBD complexes with three different classes of antibodies were used in our investigation. These structures included the SARS-CoV-2 S-RBD complexes with class I B38 antibody (Figure 1A), class II P2B-2F6 antibody (Figure 1B), and class III EY6A and S304 antibodies (Figure 1C). All structures were obtained from the Protein Data Bank.^69, 70^ Hydrogen atoms and missing residues were initially added and assigned according to the WHATIF program web interface.^71, 72^ The missing loops in the cryo-EM structures were also reconstructed using template-based loop prediction approaches ModLoop^73^ and ArchPRED.^74^ The side chain rotamers were refined and optimized by SCWRL4 tool.^75^ The protein structures were then optimized using atomic-level energy minimization with a composite physics and knowledge-based force fields using 3Drefine method.^76^ In addition to the experimentally resolved glycan residues present in the structures of studied SARS-CoV-2 S-RBD complexes, the glycosylated microenvironment was mimicked by using the structurally resolved glycan conformations for 16 out of 22 most occupied N-glycans (N122, N165, N234, N282, N331, N343, N603, N616, N657, N709, N717, N801, N1074, N1098, N1134, N1158) as determined in the cryo-EM structures of the SARS-CoV-2 spike S trimer in the closed state (K986P/V987P, pdb id 6VXX) and open state (pdb id 6VYB) and the cryo-EM structure SARS-CoV-2 spike trimer (K986P/V987P) in the open state (pdb id 6VSB).

**Figure 1.**
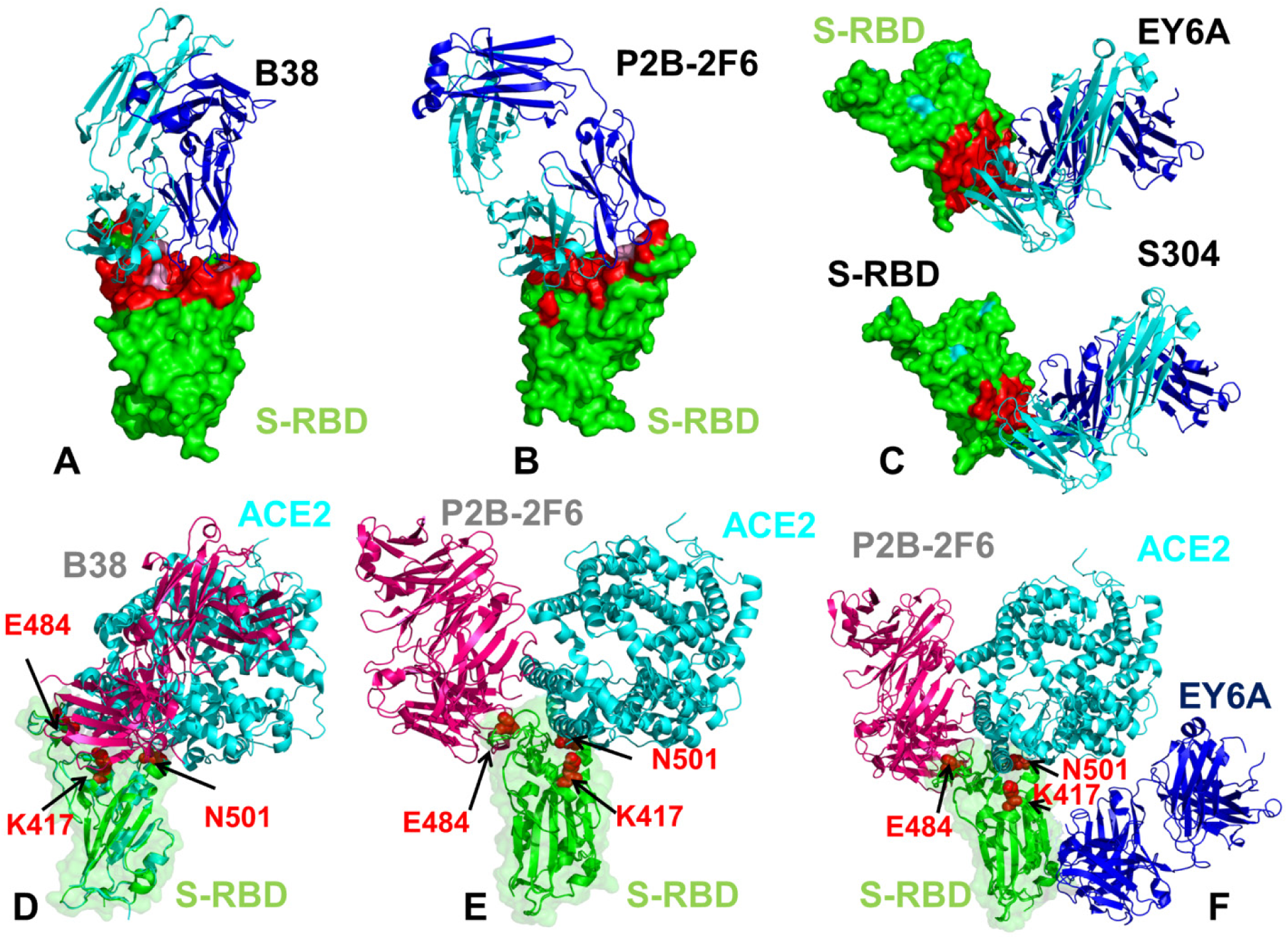
The structures of the SARS-CoV-2 S-RBD complexes with a panel of antibodies used in this study. (A) The structure of the SARS-CoV-2 S-RBD complex with B38 (pdb id 7BZ5).^19^ (B) The structure of the SARS-CoV-2 S-RBD complex with P2B-2F6 (pdb id 7BWJ).^28^ (C) The structures of the SARS-CoV-2 S-RBD complex with EY6A (pdb id 6ZER,6ZCZ)^36^ and S304 antibodies.^30^ S-RBD is shown in green surface and the binding epitopes are colored in red. The antibodies are shown in ribbons with heavy chain colored in blue and light chain in cyan. (D) Structural superposition of the B38 antibody (in pink ribbons) and ACE2 host receptor (in cyan ribbons) bound to S-RBD (in green ribbons and surface with reduced transparency). (E) Structural superposition of the P2B-2F6 antibody (in pink ribbons) and ACE2 host receptor (in cyan ribbons) bound to S-RBD (in green ribbons). (F) Structural superposition of the P2B-2F6 antibody (in pink ribbons), ACE2 host receptor (in cyan ribbons) and EY6A (in blue ribbons) bound to S-RBD (in green ribbons). The positions are sites K417, E484 and N501 subjected to circulating mutational variants are shown in red spheres and annotated.

### MD Simulations of the SARS-CoV-2 S-RBD Complexes with Antibodies

All-atom MD simulations were performed for an N, P, T ensemble in explicit solvent using NAMD 2.13 package^77^ with CHARMM36 force field.^78^ Long-range non-bonded van der Waals interactions were computed using an atom-based cutoff of 12 Å with switching van der Waals potential beginning at 10 Å. Long-range electrostatic interactions were calculated using the particle mesh Ewald method^79^ with a real space cut-off of 1.0 nm and a fourth order (cubic) interpolation. SHAKE method was used to constrain all bonds associated with hydrogen atoms. Simulations were run using a leap-frog integrator with a 2 fs integration time step. Energy minimization after addition of solvent and ions was carried out using the steepest descent method for 100,000 steps. All atoms of the complex were first restrained at their crystal structure positions with a force constant of 10 Kcal mol^-1^ Å^-2^. Equilibration was done in steps by gradually increasing the system temperature in steps of 20K starting from 10K until 310 K and at each step 1ns equilibration was done keeping a restraint of 10 Kcal mol-1 Å-2 on the protein C_α_ atoms. After the restrains on the protein atoms were removed, the system was equilibrated for additional 10 ns. An NPT production simulation was run on the equilibrated structures for 500 ns keeping the temperature at 310 K and constant pressure (1 atm). In simulations, the Nose– Hoover thermostat^80^ and isotropic Martyna–Tobias–Klein barostat^81^ were used to maintain the temperature at 310 K and pressure at 1 atm respectively. Principal component analysis (PCA) of MD trajectories using the CARMA package.^82^ We also computed the relative solvent accessibility parameter (RSA) that is defined as the ratio of the absolute solvent accessible surface area (SASA) of that residue observed in a given structure and the maximum attainable value of the solvent-exposed surface area for this residue using web server GetArea. ^83^

### Distance Fluctuations Stability Analysis

Using a protein mechanics-based approach^84, 85^ we employed distance fluctuation analysis of the simulation trajectories to profile antibody-induced modulation of stability for the SARS-CoV-2 S-RBD residues. We computed the fluctuations of the mean distance between each atom within a given residue and the atoms that belong to the remaining residues of the protein. The fluctuations of the mean distance between a given residue and all other residues in the ensemble were converted into distance fluctuation stability indexes that measure the energy cost of the residue deformation during simulations. The high values of distance fluctuation indexes are associated with stable residues that display small fluctuations in their distances to all other residues, while small values of this stability parameter would point to more flexible sites that experience large deviations of their inter-residue distances. The distance fluctuation stability index for each residue is calculated by averaging the distances between the residues over the simulation trajectory using the following expression:

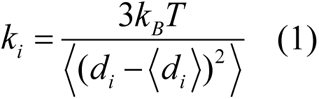

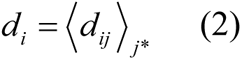

*d_ij_* is the instantaneous distance between residue *i* and residue *j*, *k_B_* is the Boltzmann constant, *T* =300K. 〈 〉 denotes an average taken over the MD simulation trajectory and 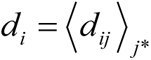 is the average distance from residue *i* to all other atoms *j* in the protein (the sum over *j*_*_ implies the exclusion of the atoms that belong to the residue *i*). The interactions between the *C_α_* atom of residue *i* and the *C_α_* atom of the neighboring residues *i* −1 and *i* +1 are excluded in the calculation since the corresponding distances are nearly constant. The inverse of these fluctuations yields an effective force constant *k_i_* that describes the ease of moving an atom with respect to the protein structure.

### Mutational Scanning

To compute protein stability changes in the SARS-CoV-2 S-RBD complexes, we conducted mutational sensitivity scanning of protein residues by systematically exploring all possible substitutions. BeAtMuSiC approach was employed that is based on statistical potentials describing the pairwise inter-residue distances, backbone torsion angles and solvent accessibilities, and considers the effect of the mutation on the strength of the interactions at the interface and on the overall stability of the complex.^86^ The binding free energy of protein-protein complex can be expressed as the difference in the folding free energy of the complex and folding free energies of the two protein binding partners:

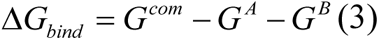

The change of the binding energy due to a mutation was calculated then as the following:

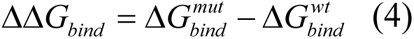

We leveraged rapid calculations based on statistical potentials to compute the ensemble-averaged binding free energy changes using equilibrium samples from MD trajectories. The binding free energy changes were computed by averaging the results over 1,000 equilibrium samples for each of the studied systems.

### Perturbation Response Scanning

Perturbation Response Scanning (PRS) approach^87–89^ follows the protocol originally proposed by Bahar and colleagues^90, 91^ and was described in detail in our previous studies.^92, 93^ In brief, through monitoring the response to forces on the protein residues, the PRS approach can quantify allosteric couplings and determine the protein response in functional movements. In this approach, it 3N × 3*N* Hessian matrix ***H*** whose elements represent second derivatives of the potential at the local minimum connect the perturbation forces to the residue displacements. The 3*N*-dimensional vector **Δ*R*** of node displacements in response to 3*N*-dimensional perturbation force follows Hooke’s law ***F* = *H \** Δ*R A*** perturbation force is applied to one residue at a time, and the response of the protein system is measured by the displacement vector Δ***R***(*i*) = ***H*^-1^F**^(*i*)^ that is then translated into *N*×*N* PRS matrix. The second derivatives matrix ***H*** is obtained from simulation trajectories for each protein structure, with residues represented by *C*_*α*_ atoms and the deviation of each residue from an average structure was calculated by Δ**R**_*j*_(*t*) = **R**_*j*_(*t*) −〈**R**_*j*_(*t*)〉, and corresponding covariance matrix C was then calculated by Δ**R**Δ**R**^*T*^. We sequentially perturbed each residue in the SARS-CoV-2 spike structures by applying a total of 250 random forces to each residue to mimic a sphere of randomly selected directions.^62^ The displacement changes, Δ***R***^***i***^ is a *3N-*dimensional vector describing the linear response of the protein and deformation of all the residues.

Using the residue displacements upon multiple external force perturbations, we compute the magnitude of the response of residue *k* as 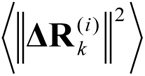 averaged over multiple perturbation forces **F**^(^*^i^*^)^, yielding the *ik*^th^ element of the *N*×*N* PRS matrix. The average effect of the perturbed effector site *i* on all other residues is computed by averaging over all sensors (receivers) residues *j* and can be expressed as〈(Δ***R***^***i***^)^2^〉*_effector_*. The effector profile determines the global influence of a given residue node on the perturbations in other protein residues and can be used as proxy for detecting allosteric regulatory hotspots in the interaction networks. In turn, the *j*^th^ column of the PRS matrix describes the sensitivity profile of sensor residue *j* in response to perturbations of all residues and its average is denoted as 〈(Δ***R***^***i***^)^2^〉*_sensor_*. The sensor profile measures the ability of residue *j* to serve as a receiver of dynamic changes in the system.

## Results and Discussion

### Dynamic Signatures of the SARS-CoV-2 S RBD Are Uniquely Modulated by Different Classes of Antibodies

Using MD simulations, we examined how SARS-CoV-2 spike protein can exploit plasticity of the RBD regions to modulate specific dynamic responses to antibody binding. MD simulations of the RBD-antibody complexes revealed functionally important modulation of the conformational dynamics and appreciable redistribution of the S-RBD stability profiles. The RBD core remained stable during while mobility of some flexible regions could be significant altered (Figure 2). Instructively, the largest binding-induced dynamic changes were seen in the S-RBD complex with B38, exemplified by significant stabilization of flexible residues around K417 site (Figure 2A). An appreciable modulation of the S-RBD mobility was also seen in the loop region (residues 457-480) and RBM ridge (residues 490-505) that display moderate thermal fluctuations in the complex (Figure 2A). The strategic functional positions K417 and N501 are stabilized in the RBD-B38 complex, but residues around E484 position are noticeably more flexible (Figure 2A). The distribution of the distinct inter-molecular residue pairs revealed antibody contacts with a wide range of residues including D420, K417, Y421, K458, G476, D427, F486, Y489, Q498, N501, Y505 of the SARS-CoV-2 RBD (Supporting Information, Figure S1A). In particular, K417 forms hydrophilic interactions with Y33, N92, Y52, Y58 and Y97 of B38. Another important site N501 is involved in multiple interactions with various B38 residues of the light chain (Q27, S30, G28,Q 27, I29). Notably, most interactions at the RBD-B38 interface are hydrophilic formed by K417, D403, D420, K458, and Q498 residues. A number of the inter-residue contact pairs are mediated through interactions of hydrophobic hotspots F456, F486, and Y489 (Supporting Information, Figure S1A). A similar dynamic pattern was detected in the S-RBD complex with class II P2B-2F6 antibody that binds to the RBM region using a different recognition angle with the binding epitope centered on G446/Y449 and E484/F490 residues (Figure 2B). These key epitope residues showed modest thermal fluctuations and were stabilized in the complex, but the overall mobility of the S-RBD was greater as compared the RBD-B38 complex. The inter-residue interaction pairs are formed with V445, G446, and Y449 site (Supporting Information, Figure S2B). In particular Y449 residue is involved in numerous contacts with Y27, V104, G32, S31, Y33, and G102 of heavy chain of B38. Of special notice also a number of the inter-residue pair contacts made by E484, F486, and F490 residues (Supporting Information, Figure S1B). E484 forms hydrogen bonds with N33 and Y34 of the P2B-2F6 light chain and with R112 of the P2B-2F6 heavy chain.

**Figure 2.**
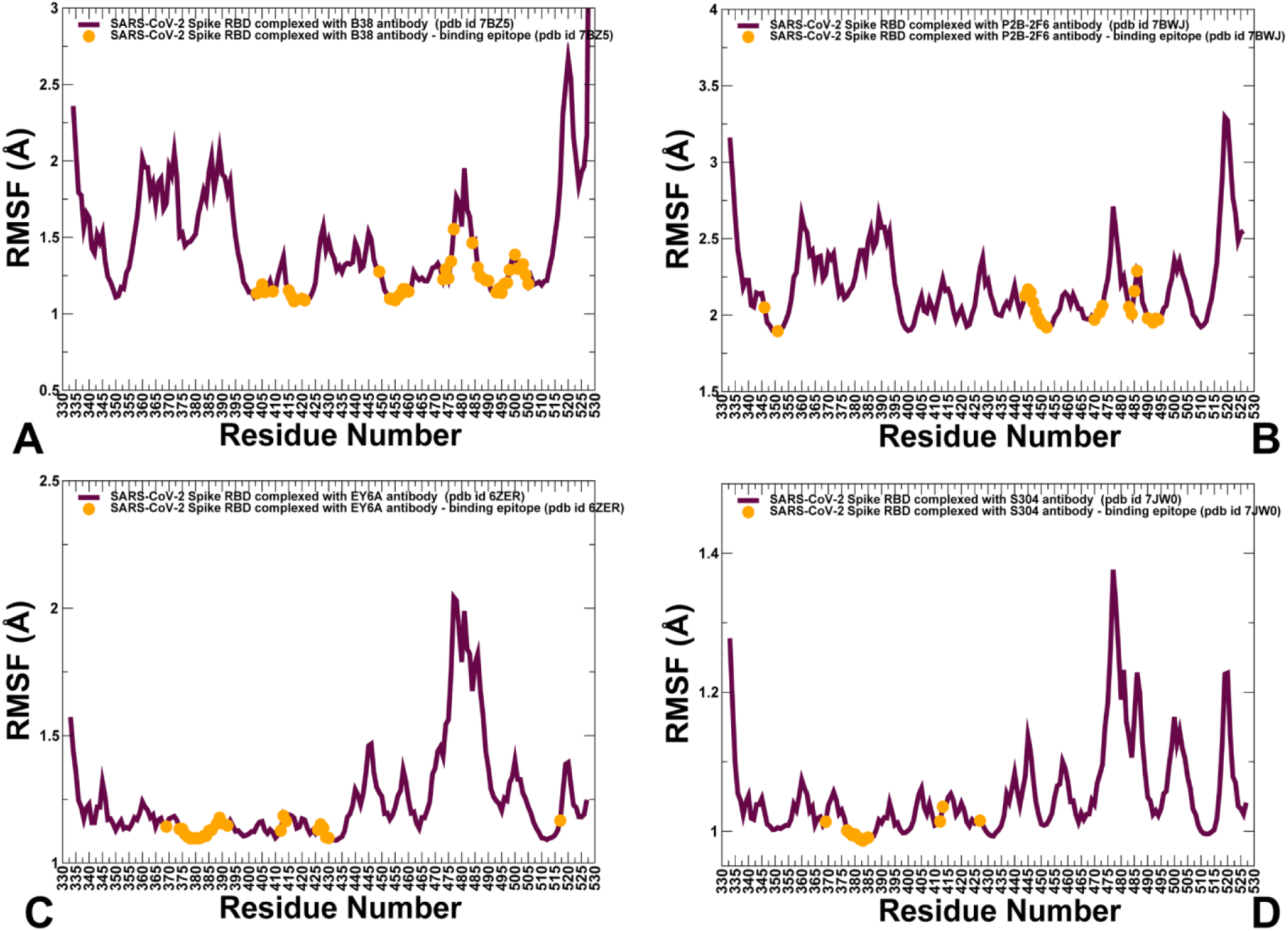
Conformational dynamics of the SARS-CoV-2 S-RBD complexes with different classes of antibodies. (A) The root mean square fluctuations (RMSF) profiles from MD simulations of the structures of the SARS-CoV-2 S-RBD complex with class I B38 antibody (pdb id 7BZ5, shown in maroon lines). (B) The RMSF profiles from MD simulations of the structures of the SARS-CoV-2 S-RBD complex with class II P2B-2F6 antibody (pdb id 7BWJ). (C,D) The RMSF profiles from MD simulations of the structures of the SARS-CoV-2 S-RBD complex with class III EY6A and S304 antibodies. The binding epitope residues in the S-RBD complexes with antibodies are highlighted in orange filled circles.

The class III antibodies EY6A and S304 bind to the cryptic epitope and exhibited a characteristic dynamics profile showing a considerable stabilization of the antibody-interacting residues and further rigidification of the RBD core (Figure 2C,D). In comparison, one moderately flexible region corresponded to the 444–451 and 456-460 loops, and another peak of the distribution was in the exposed region (residues 480-486) from the tip of the RBM loop (Figure 2C,D).

In this context, it is worth noting that mutations in the epitope centered around E484 position (G485, F486, F490) or in the 443–455 loop (K444, V445, L455, F456) can strongly affect neutralization and trigger resistance for various classes of antibodies, but are less detrimental for binding of EY6A and S304 antibodies where such mutations may be more readily accommodated.^36^ Notably, this class of antibodies may not only promote further stabilization of the RBD core but could also modulate mobility of the RBM region. As a result, even though EY6A binding does not impose steric hindrance on interactions with ACE2 (Figure 1F), by differentially modulating stability of the RBM residues EY6A may allosterically induce a weak inhibitory effect on binding with the host receptor.^36^

A similar distribution of the inter-residue pairs can be mediated by EY6A and S304 antibodies. For EY6A, the large numbers of contacts are established by G381, V382, S383, P384, T385, and K386 residues that form the core of the binding epitope (Supporting Information, Figure S1C). A less dense network of contacts is maintained by S304 antibody in the complex, featuring C379, G381, V382, and S383 RBD residues forming contacts with distinct positions on both heavy and light chains (Supporting Information, Figure S1D).

We compared profiles for the unbound SARS-CoV-2 S-RBD form with the respective distributions in the antibody-bound states to determine how binding modulates conformational mobility of the S-RBD residues. Using the ensemble-based distance fluctuations analysis of the unbound SARS-CoV-2 S-RBD we first identified intrinsically structurally stable regions and characterize dynamic signatures of the binding epitopes (Supporting Information, Figure S2). The analysis highlighted several important “islands” of structural stability that are primarily located in the RBD core. The distribution showed clear peaks corresponding to the conserved RBD core consisting of antiparallel β strands (β1 to β4 and β7) (residues 354-358, 376-380, 394-403, 431-438, 507-516). Noticeably, the β-sheets β5 and β6 (residues 451-454 and 492-495) that anchor the RBM region to the central core also featured high values of the stability index. A number of structurally stable RBD residues corresponded to evolutionary conserved positions including C336, R355, C361, F374, F377, C379, L387, C391, D398, G413, N422, Y423, L425, F429, C432, and W436 (Supporting Information, Figure S2). The distance fluctuation stability profile suggested a moderate stability of K417 site, while E484 and N501 positions featured very small index values indicative of high mobility in the free form. By mapping the binding epitope residues for studied antibodies onto distance fluctuation profile of the S-RBD, we evaluated the intrinsic dynamic preferences of the binding interfaces (Supporting Information, Figure S2). The large binding epitope for B38 featured stable segments (residues 403-421, 453-460) as well as more flexible region (residues 473-480) (Supporting Information, Figure S2A). Interestingly, the binding epitopes for EY6A and S304 antibodies targeting the conserved RBD core included moderately stable RBD residues, and only residues F377 and F429 were aligned with the local maxima of the profile (Supporting Information, Figure S2C,D).

The distance fluctuation stability profiles for the S-RBD complexes with antibodies revealed how binding can induce changes in the distribution of rigid and flexible regions (Figure 3). In the complex with B38 antibody, one could notice a significant stabilization of the RBM residues involved in the binding interactions (Figure 3A). In particular, functional sites K417 and N501 become considerably more stable while E484 residue retained relatively high mobility present in the unbound RBD form. Despite the overlap of the binding epitopes for B38 and P2B-2F6 antibodies that target the RBM region, the stability profiles for these complexes featured several specific signatures (Figure 3B). The distance fluctuation profile for the P2B-2F6 complex pointed to a moderate mobility of K417 and N501 sites, while revealing a markedly increased stabilization of highly mobile region centered on E484 in the RBM region (Figure 3B). The distributions for class III EY6A and S304 antibodies were quite similar due to very similar binding mode and interactions targeting the cryptic binding site (Figure 3C,D). Of particular notice are pronounced and sharp peaks associated with the conserved residues in the RBD core that become largely rigid in the complexes. On the other hand, these antibodies induced a more significant mobility in the RBM regions (residues 470-490). Notably, functional sites K417, E484 and N501 displayed relatively low values of the stability index (Figure 3C,D). In addition, we noticed a more radical segregation of highly stable and flexible regions in these complexes where the rigidity of the RBD core residues can be contrasted with mobility of the RBM residues. Overall, this analysis indicated that different classes of antibodies could differentially modulate stability of the S-RBD residues. Moreover, binding of these antibodies can uniquely specify dynamic signatures of functional regions targeted by global mutational variants.

**Figure 3.**
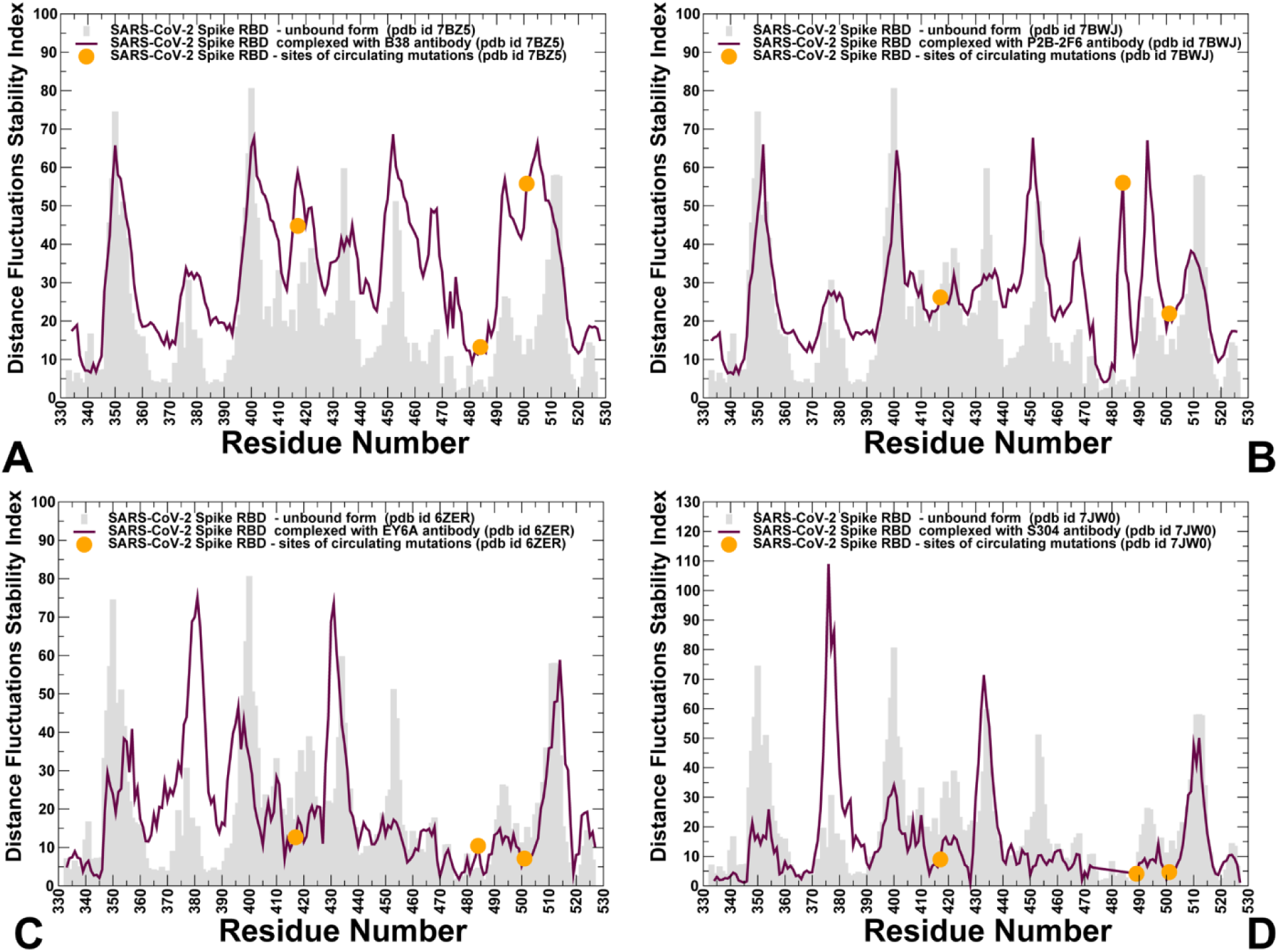
The distance fluctuations analysis of the conformational ensembles in the SARS-CoV-2 S-RBD complexes with different classes of antibodies. (A) The distance fluctuation stability index for the SARS-CoV-2 S-RBD complex with class I B38 antibody (pdb id 7BZ5). (B) The distance fluctuation stability index for the SARS-CoV-2 S-RBD complex with P2B-2F6 antibody (pdb id 7BWJ). (C,D) The distance fluctuation stability index for the SARS-CoV-2 S-RBD complexes with class III EY6A and S304 antibodies. The stability index profile for the unbound form of the S-RBD is shown on panels (A-D) in light grey bars. The stability index profiles for the SARS-CoV-2 S-RBD complexes are shown in maroon-colored lines. The position of functional RBD sites K417, E484, and N501 targeted by global circulating variants are highlighted in orange filled circles.

We specifically mapped stable spike residues that form local modular clusters in the structures of the SARS-CoV-2 S-RBD complexes with antibodies (Figure 4). These structurally stable residues in the complex with class I B38 antibody connect the peripheral spike regions through the RBD core to the binding interface (Figure 4A). Moreover, the functional sites K417, N501 and E484 appeared to form a network of intermolecular bridges that link the stable RBD core with the antibody. Interestingly, although P2B-2F6 interactions with S-RBD are centered on E484 site, these interactions are fairly isolated and separated from the dense clusters of stable residues of the RBD core (Figure 4B). In this complex, N501 residue provided the link between stable RBD clusters and the inter-molecular interface. For class III antibody EY6A, the binding interface is linked with the stable RBD core, illustrating the observed rigidification of these regions (Figure 4C). Functional centers of K417 and N501 are directly linked to these stable clusters suggesting that these positions could sense the antibody-induced signal and alter their dynamics as a part of protein response.

**Figure 4.**
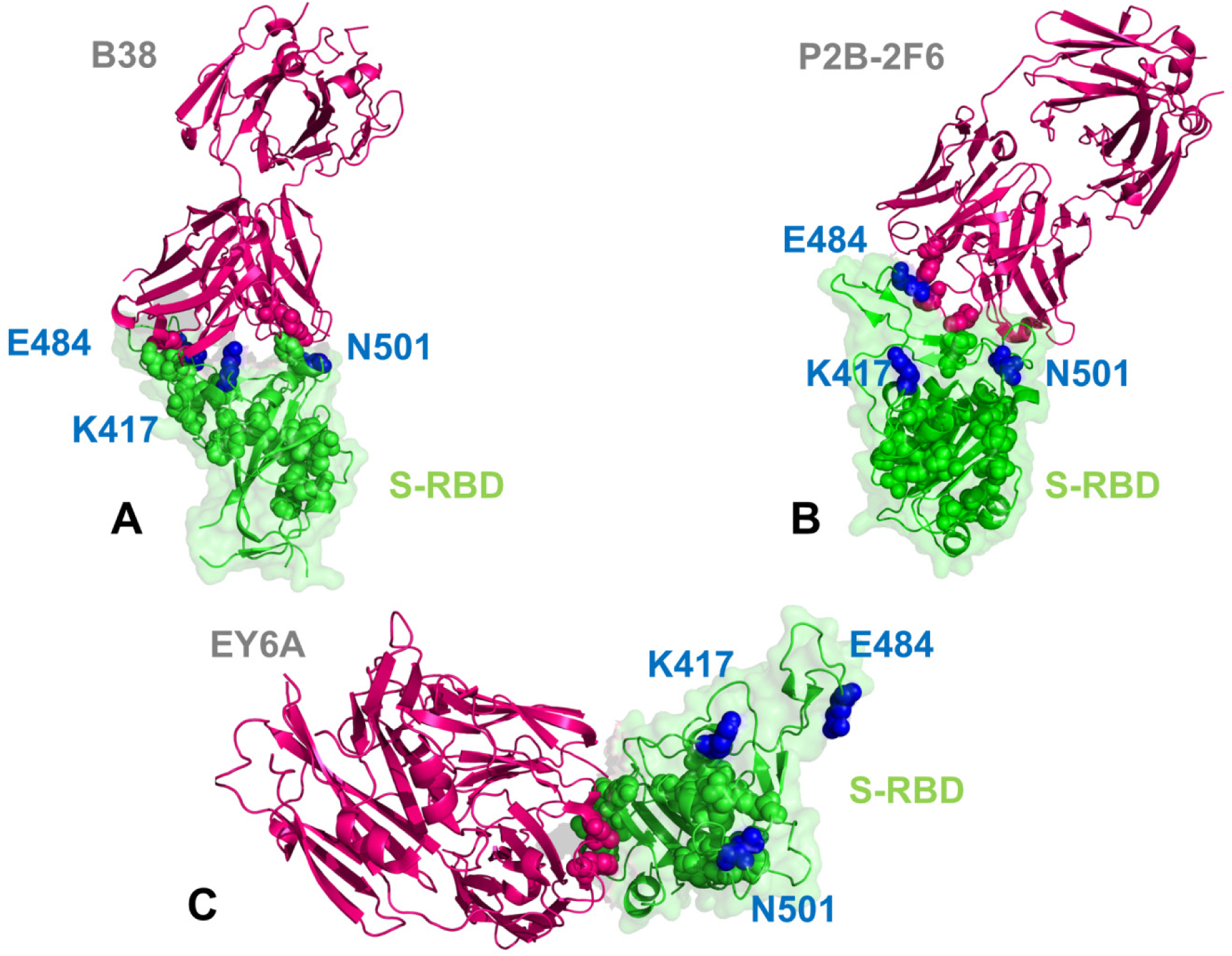
Structural mapping of local clusters of stable RBD residues revealed by the distance fluctuation analysis. Structural analysis of the SARS-CoV-2 S-RBD complexes with class I B38 antibody (A), class II P2B-2F6 antibody (B) and class III EY6A antibody (C). The S-RBD is shown in green ribbons and surface representation with a reduced transparency. The antibodies on panels (A-D) are shown in dark-pink ribbons. The stable RBD residues forming local clusters are show in green spheres. The functional RBD sites K417, E484, and N501 targeted by global circulating variants are shown in blue spheres and annotated.

To enhance this dynamics-based analysis, we also employed the conformational ensemble to compute the relative solvent accessibility (RSA) ratio in the SARS-CoV-2 RBD complexes. This parameter was obtained by averaging the SASA computations over the simulation trajectories (Figure 5). Of particular interest were the average RSA values for functional sites K417, E484, and N501 that are residues targeted by circulating mutations. Interestingly, B38 and P2B-2F6 antibodies yielded a different pattern of RSA for these sites. For B38-RBD complex, we observed that K417 and N501 positions are buried featuring RSA < 10% (Figure 5A), while E484 position is largely solvent-exposed. In the complex with P2B-2F6, both N501 and especially E484 become > 80% buried (RSA< 20%) while K417 site remained partly exposed to solvent (Figure 5B).

**Figure 5.**
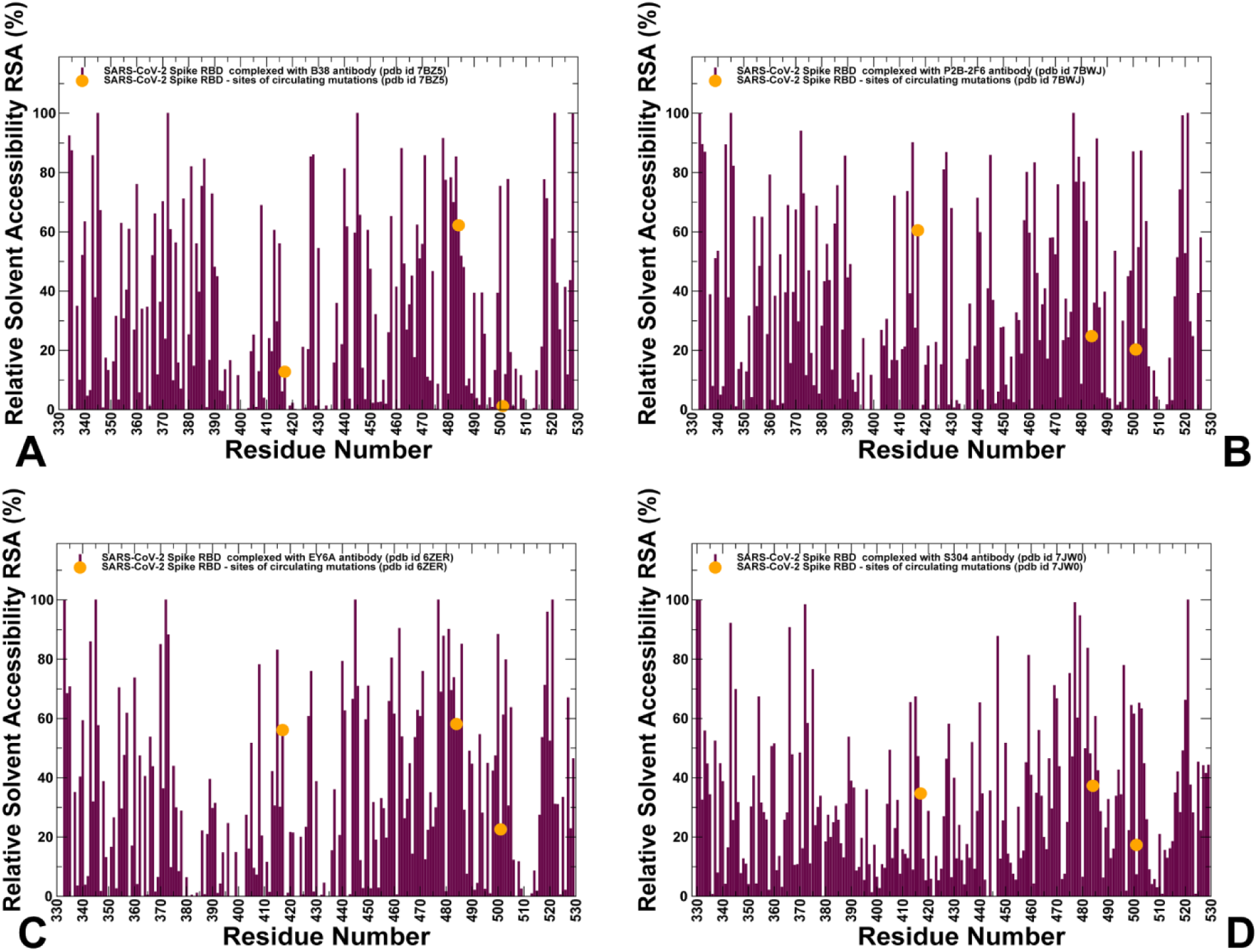
The average relative solvent accessibility (RSA) percentage of the protein residues for the SARS-CoV-2 S-RBD complexes with different classes of antibodies. (A) The RSA profile for the SARS-CoV-2 S-RBD complex with class I B38 antibody (pdb id 7BZ5). (B) The RSA profile for the SARS-CoV-2 S-RBD complex with P2B-2F6 antibody (pdb id 7BWJ). (C,D) The RSA profile for the SARS-CoV-2 S-RBD complexes with class III EY6A and S304 antibodies. The RSA profiles are shown in maroon lines and positions of functional RBD sites K417, E484, and N501 targeted by global circulating variants are highlighted in orange filled circles.

This class of antibodies revealed key interactions with residue E484, showing most of these antibodies can be evaded by 501Y.V2 RBD variant that prominently featured E484K mutation.^27^ Because EY6A and S304 antibodies target the cryptic site located away from the RBD regions, the functional sites K417 and E484 were mostly exposed with N501 only moderately buried (Figure 5C,D). It should be pointed out that surrounding residues in this RBM ridge (T500, V503, G504, and T505) are completely exposed featuring RSA values between 65% and 80%. Hence, different classes of antibodies may induce distinct patterns of solvent accessibility in the functional positions subjected to circulating mutations.

### Binding-Induced Modulation of Soft Collective Modes Determines Unique Protein Responses to Different Antibody Classes

We also characterized collective motions for the SARS-CoV-2 S-RBD complexes averaged over low frequency modes using principal component analysis (PCA) of the MD trajectories (Figure 6). It is worth noting that the local minima along these profiles are typically aligned with the immobilized in global motions hinge centers, while the maxima correspond to the moving regions undergoing concerted movements leading to global changes in structure.^95^ The low-frequency ‘soft modes’ are characterized by their cooperativity, functional significance and robustness whereby there is a strong relationship between allosterically-driven conformational changes and the ‘soft’ modes of motions intrinsically accessible to folded structures.^95.96^ It is well established that allosteric responses in proteins can be efficiently triggered when external stimuli (such as mutations, ligand or antibody binding) can exploit the intrinsic protein propensities for energetically favorable movement along the slow modes.^96^ In general, allosteric effects often arise when the pre-existing slow modes can be altered upon ligand binding. As a result, when antibody binding to the S-RBD protein target flexible regions near E484 and N501 positions, this may elicit functionally relevant and cooperative allosteric response and change functional motions of the spike protein. Our analysis is particularly instructive when the slow mode profiles of the S-RBD are compared between the unbound and bound spike forms. In the unbound S-RBD form the profile featured a very pronounced density peak centered on E484/F486 residues and a next peak around position N501 (Figure 6). At the same time, another functional site K417 was located near a relatively shallow maximum along this profile. This suggested that sites of circulating mutations belong to the functional regions undergoing concerted motions along pre-existing slow modes of the spike protein. Interestingly, none of the functional positions targeted by novel mutational variants that promote infectivity and antibody resistance (K417, E484, and N501) corresponded to hinge positions in the unbound RBD form. Hence, targeting these regions by antibodies could potentially change the spectrum of slow modes and induce specific functional motions characteristic for different antibody classes.

**Figure 6.**
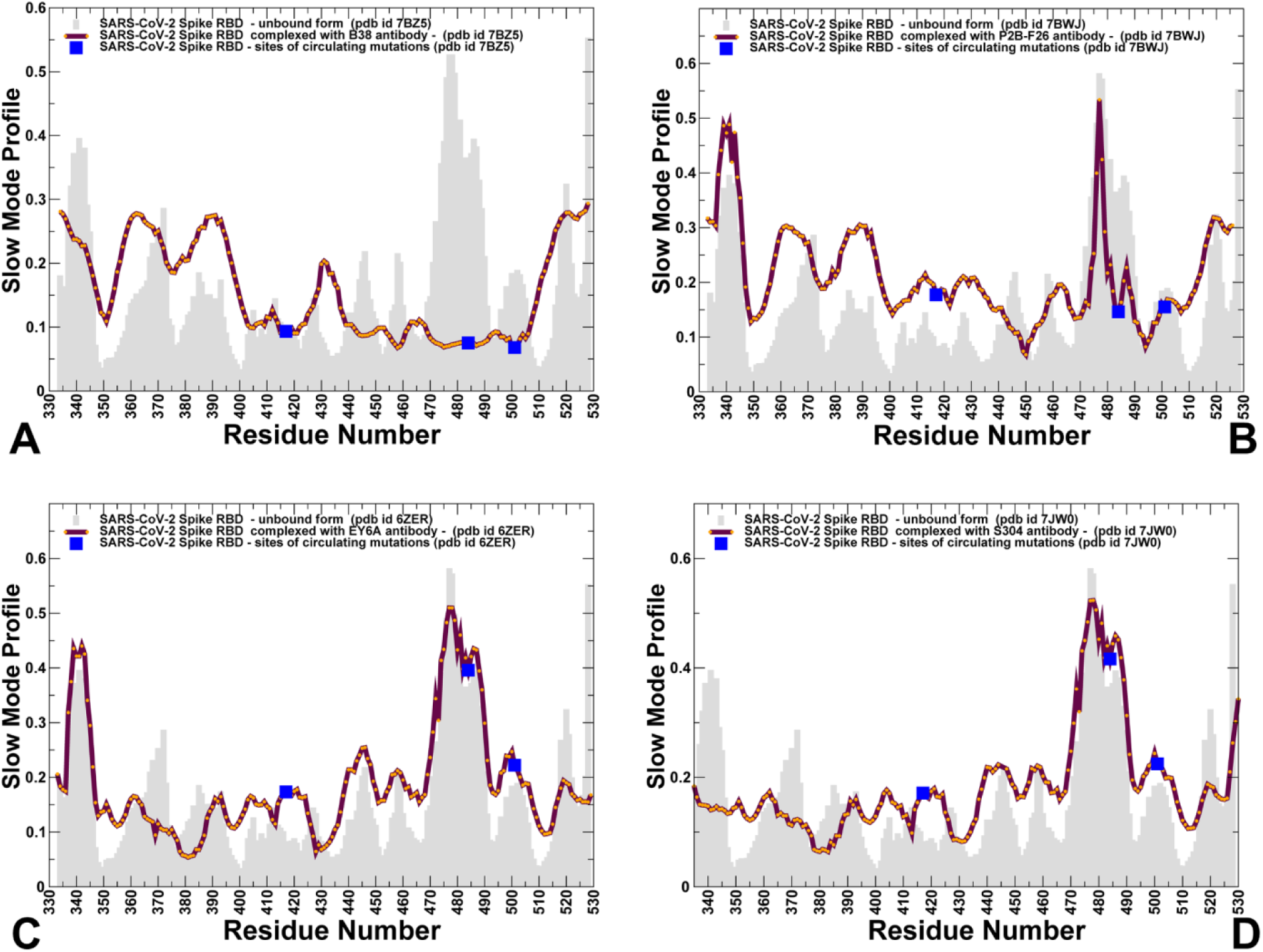
Collective dynamics of the SARS-CoV-2 S-RBD complexes with different classes of antibodies. The mean square displacements in functional motions are averaged over the three lowest frequency modes. (A) The essential mobility profile for the SARS-CoV-2 S-RBD complex with class I B38 antibody (pdb id 7BZ5). (B) The slow mode profile for the SARS-CoV-2 S-RBD complex with P2B-2F6 antibody (pdb id 7BWJ). (C,D) The essential mobility distribution for the SARS-CoV-2 S-RBD complexes with class III EY6A and S304 antibodies. The slow mode profile for the unbound form of the S-RBD is shown on panels (A-D) in light grey bars. The slow mode profiles for the SARS-CoV-2 S-RBD complexes are shown in maroon-colored lines with individual data points highlighted in orange circles. The position of functional RBD sites K417, E484, and N501 targeted by global circulating variants are highlighted in filled blue squares.

Indeed, for the class I B38 antibody whose binding mode is centered on K417 region, the global mobility distribution was markedly altered, showing how binding can shifty the location of major hinge sites (Figure 6A). We noted that B38 binding did not affect hinge residues in the RBD core as residues Y351, F374, and G404 were aligned with the local minima of the profile. At the same time, the slow mode profile for the complex revealed local minima near K417, E484 and N501 residues (Figure 6A). Moreover, the entire moving region of the unbound from (residues 472-486) becomes mostly restricted in the global dynamics of the complexed SARS-CoV-2 RBD (Figure 6A). This indicated a substantial redistribution of stable hinge site positions where all three sites targeted by circulating mutations become recruited to the pool of hinge sites and regulatory centers in the S-RBD protein. As a result, these functional positions may be involved in a cooperative cross-talk with each other and other conserved hinges located away from the ACE2 binding site. We argue that B38-induced changes in collective dynamics could strengthen both binding interactions and allosteric communications in the complex. On the flipped side, mutations in the triad K417/E484/N401 may not only affect local binding interactions but would also alter the allosteric response of moving positions that are sensitive to the specific perturbation, leading to changes in global dynamics and compromising antibody-induced allosteric response. In this context it is particularly interesting to compare our observations with functional studies showing that E484K/N501Y/K417N mutations in the 501Y.V2 lineage confer neutralization escape from class I of SARS-CoV-2 antibodies.^27^

In the complex with class II antibody P2B-2F6, the hinge sites are aligned with Y449, S469, E484 and S494 residues (Figure 6B). A deep global minimum is centered on Y449 residue which is the key position of the P2B-2F6 binding epitope within the crevice formed by the RBM-hairpin. Moreover, E484 residue is now aligned with a local hinge position and K417 belongs to a shallow minimum, while N501 peak is aligned with the respective peak in the unbound state (Figure 6B). Our analysis indicated that G446, Y449 and E484 anchoring positions are critical for functional motions in the complex and mutations in these sites could alter binding to P2B-2F6 antibody. Unlike sites of circulating mutations, modifications in positions G446, Y4459 could compromise binding of SARS-CoV-2 RBD with both ACE2 and the antibody. At the same time, mutations in E484 would likely have severe effect on antibody recognition and functional motions.

A completely different pattern was seen in the collective dynamics profiles of SARS-CoV-2 RBD complexes with EY6A and S304 antibodies targeting the cryptic site (Figure 6C,D). The hinge centers in the RBD core that are intrinsic for the unbound form were prominently present in the profiles of these complexes, particularly pointing to cluster of residues 381-386 and 427-432. These stable RBD core positions become immobilized due to additional stabilizing contacts and form the regulatory control center of functional movements in the complexes (Figure 6C,D). Interestingly, the slow mode profiles for these complexes are very similar to the distributions seen for the unbound S-RBD, suggesting that binding to the cryptic site may only moderately affect functional motions of the spike protein. Furthermore, the positions of functional sites K417, E484 and N501 on the slow mode profiles remained similar in the unbound and bound forms aligned with the moving regions in collective dynamics (Figure 6C,D). Moreover, binding of these may preserve or amplify mobility of the key regions surrounding E484 and N501 residues. This may allow for simultaneous binding of ACE2 albeit prone to the experimentally detected faster dynamic exchange and dissociation.^36^ According to the experiments, attachment of preincubated RBD and EY6A to immobilized ACE2 indicated that the off-rate from ACE2 was increased by the presence of EY6A, suggesting a crosstalk between the binding of ACE2 and EY6A antibodies.^36^ Our analysis indicated that EY6A and S304 antibodies could potentially act as modulators of the conformational dynamics without dramatically altering collective motions and allosteric protein response but rather fine-tune dynamic changes at the ACE2-binding site. In this model, antibody binding would strengthen allosteric couplings between ACE2-binding site and cryptic binding site, arguably facilitating a cross-talk between bound ACE2 and antibodies.

### Mutational Scanning Heatmaps Identify Binding Energy Hotspots and Interaction Propensities of Functional Sites in the SARS-CoV-2 Complexes with Antibodies

To provide further comparison between the computational and experimental data, we performed mutational sensitivity analysis and constructed mutational heatmaps for SARS-CoV-2 S-RBD binding (Figure 7). The binding epitope of the class I antibody B38 has a considerable overlap with the ACE2 binding site (Figure 1). In fact, 21 residues of the SARS-CoV-2 S-RBD are shared between epitope residues bound by B38 and ACE2. The unique epitope residues that interact with the B38 antibody include D405, E406, T415, G416, D420, Y421, N460, Y473, A475, E484, and G496 (Figure 7A). However, only several of these positions (Y421, Y473, A475 residues) corresponded to the binding hotspots (Figure 7A). The computed mutational scanning map for the S-RBD binding with B38 pointed to sites K417, Y421, Y453, F456, N487, Q493, and Y505 as the most dominant binding energy hotspots, displaying large destabilization loss in these positions for all substitutions (Figure 7A). These results are in good agreement with deep mutational scanning analysis.^19, 42, 45^ A considerably smaller binding footprint was found for class II antibody P2B-2F6 binding where major binding hotspots corresponded to the residue cluster surrounding key anchoring position of Y449 (Figure 7B). All mutations in this position produced large destabilization changes. A similar mutational scanning signature was observed in positions G447, N450, L452, V483, E484, G485, and F490 (Figure 7B). Notably, the most prominent binding energy hotspots corresponded to G447, Y449, E484, and F490 residues. The computed mutational maps reproduced the experimental data by identifying important epitope residues in the cryptic binding site targeted by EY6A and S304 (Figure 7C).

**Figure 7.**
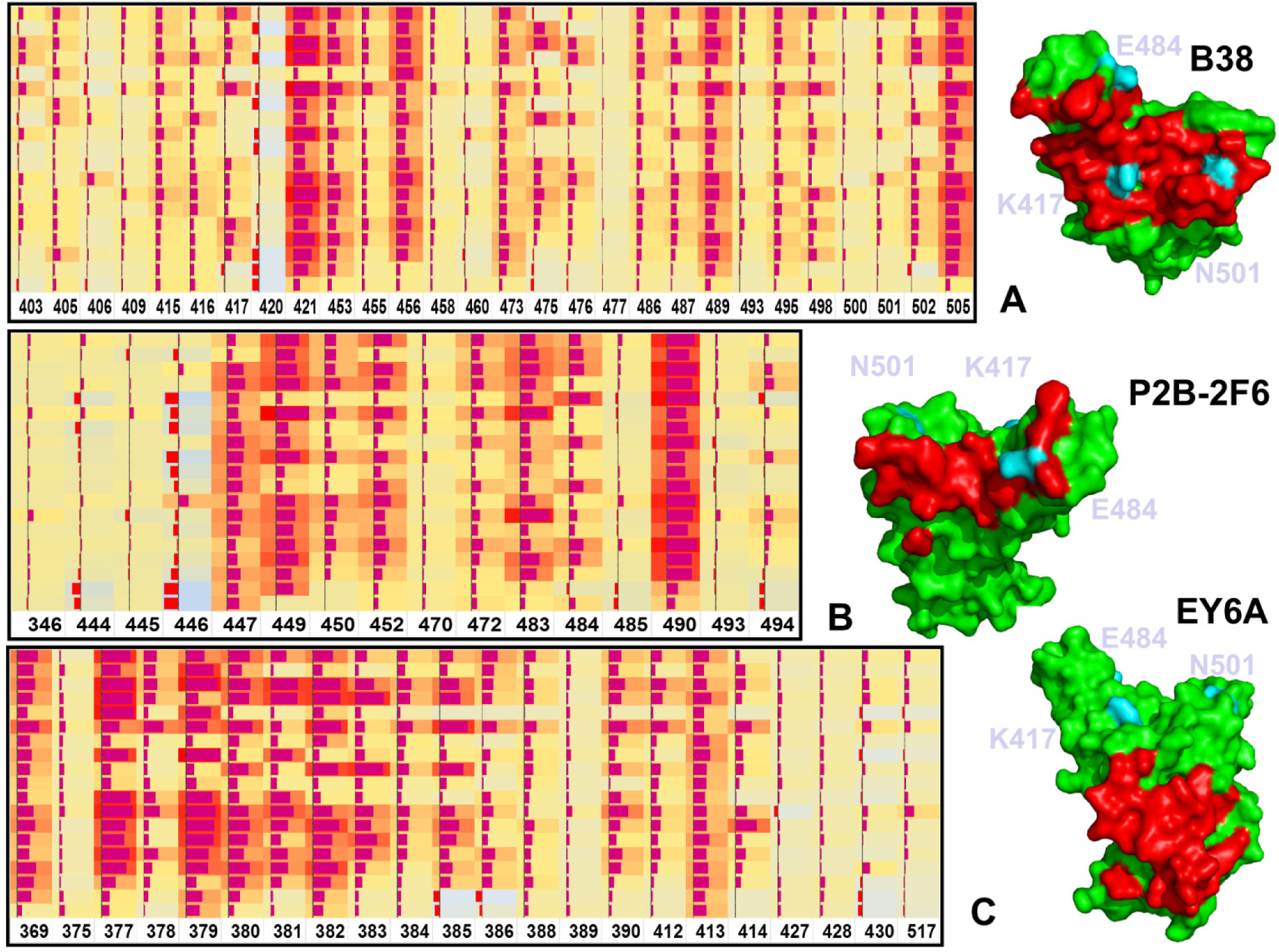
The mutational scanning heatmaps for the SARS-CoV-2 S-RBD complexes with class I B38 antibody (A), class II P2B-2F6 antibody (B) and class III EY6A antibody (C). The heatmaps illustrate the computed average binding free energy effect of all 19 single mutations on the binding epitope sites. The squares on the heatmap are colored by the mutational effect using a 3-colored scale - from light blue to red, with red indicating the largest destabilization effect. The data bars correspond to the computed binding free energy changes, where positive values (destabilizing mutations) are shown by bars towards the right end of the cell and negative values (stabilizing mutations) are shown as bars oriented towards left end. The length of the data bar is proportional to the value in the square cell. The heatmaps are shown alongside of the S-RBD structures highlighting the antibody epitopes (right side of each panel). The S-RBD is shown in green surface. The binding epitope regions are shown in red. The positions of functional sites K417, E484 and N501 are shown in light cyan color and annotated.

Mutational scanning of EY6A binding with the S-RBD revealed a dense cluster of binding energy hotspots corresponding to Y369, F377, K378, C379, Y380, G381, V382, S383, P384, T385, K386 and F390 residues that form the core of the binding epitope (Figure 7C). Several other hotspots were aligned with positions P412, G413, Q414. Class III antibodies EY6A and S304 bind to the same cryptic site and share very similar binding epitopes with CR3022. In silico mutational scanning is consistent with complete mapping of antibody-escaping RBD mutations for this class where antibody recognition can be evaded by mutations of the RBD core residues F374, G381, V382, S383, and F392.^42, 45^

Despite a number of local interaction contacts formed by B38 with functional sites K417, E484 and N501 only K417 featured as an appreciable energetic hotspot. Comparison of predicted ΔΔGs shows that a number of mutations at K417 site, particularly K417A/G/D/T resulted in a large destabilization effect (Figure 8A). In contrast, very minor changes are triggered by modifications of E484 where some mutations such as E484F and E484W may yield moderate stabilization largely due to the enhanced structural stability of this flexible region (Figure 8B). Mutations of N501 residues are generally moderately detrimental to binding with B38 with the exception of N501A and N501Y modifications that caused a more significant loss of binding (Figure 8C). We found that circulating mutations K417N (Figure 8A) and N501Y (Figure 8C) could compromise S-RBD binding to the B38 antibody, producing the destabilization effect that can be compounded by simultaneous substitutions in these positions.

**Figure 8.**
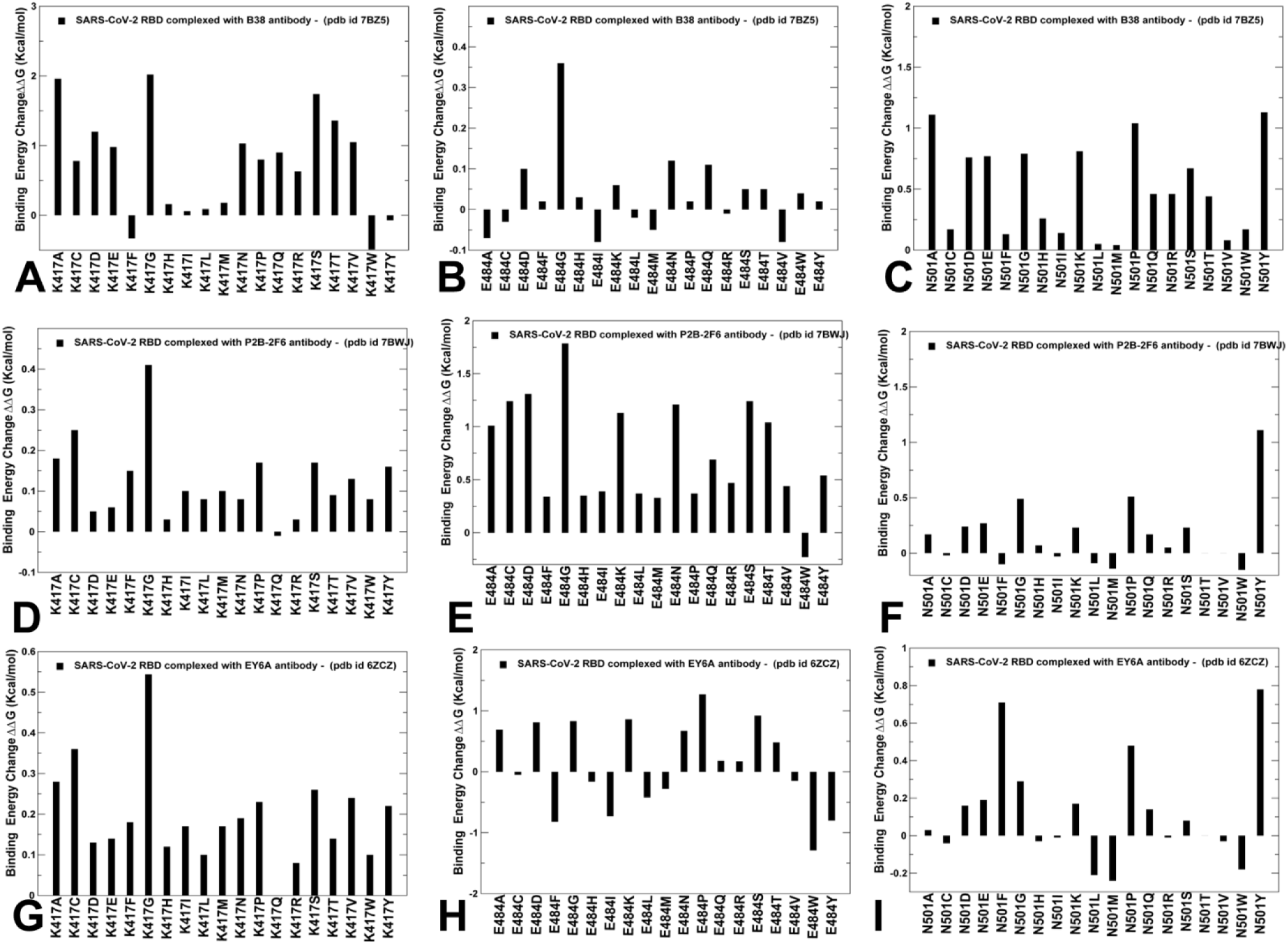
Mutational scanning of functional centers in the SARS-CoV-2 S-RBD complexes with antibodies. Mutational sensitivity scanning of K417 site (A), E484 site (B) and N501 residues (C) in the SARS-CoV-2 S-RBD complex with B38. Mutational sensitivity scanning in the SARS-CoV-2 S-RBD complex with P2B-2F6 antibody for K417 site (D), E484 site (E) and N501 residues (F). Mutational sensitivity scanning in the SARS-CoV-2 S-RBD complex with EY6A antibody for K417 site (G), E484 site (H) and N501 residues (I). The protein stability changes are shown in maroon-colored filled bars.

The results of in silico mutational screening that are based on knowledge-based mean force potentials are consistent with more detailed and laborious quantum chemical-based fragment molecular orbital (FMO) evaluation of the SARS-CoV-2 S-protein binding with the B38 Fab antibody.^97, 98^ By analyzing FMO-based interaction energies^98^, a range of residues critical for molecular recognition with B38 Fab antibody was identified that also included R403, Q409, K458, G476, N501, G502, V503, G504, and Y505 sites.

In some contrast, K417 and N501 positions are not involved in significant contacts with P2B-2F6 and located on fringes of the binding epitope. The computed free energy values upon mutations in these positions largely reflected changes in the protein stability that are very small and generally destabilizing (Figure 8D,F). Despite somewhat less dramatic destabilization changes, all mutations in E484 position showed a consistent and appreciable loss of binding affinity, including the E484K circulating mutation (Figure 8E). This implied that binding for this class of antibodies could be highly sensitive to mutations in E484 position and trigger resistance to the E484K variant. This is consistent with functional studies showing that E484K/N501Y/K417N mutations in the 501Y.V2 lineage confer a strong neutralization escape from classes I and II of SARS-CoV-2 antibodies.^27^ The results also suggested that antibodies from class I (B38) and II (P2B-2F6) can be vulnerable to distinct and non-overlapping variants as confirmed in biochemical studies^99^ According to the latest experiments, the B.1.351 (501Y.V2) variant that emerged in South Africa is resistant to neutralization by multiple monoclonal antibodies targeting the RBM of the S-RBD which is mostly due to a mutation causing an E484K substitution.^99^ These functional studies indicated that E484K mutation could trigger antibody escape for class I and II antibodies. By combining mutational scanning analysis with the results of functional dynamics profiling, one could notice that mutations in the E484 position may alter allosteric protein response for both classes of antibodies, even though binding affinity losses appeared to be significant only for the class II antibodies. This highlighted the relevance of both local and global factors induced by the mutational variants on antibody binding, where the ultimate functional response may be often affected by a non-trivial combination of binding and allosteric effects.

Functional sites K417, E484 and N501 positions are located outside of the cryptic epitope and not involved in any interactions with EY6A and S304 antibodies (Figure 7C). Mutational sensitivity profiles in these positions in the complexes with EY6A (Figure 8G-I) reflected minor changes associated with protein stability variations. Interestingly, some of these changes corresponded to the improved stability of the S-RBD, particularly for mutations in E484 and N501 sites. This indicated that functional sites E484 and N501 retain a considerable degree of structural and energetic plasticity in complexes with EY6A and S304, suggesting that circulating mutations in these positions would not confer escape from class III of SARS-CoV-2 antibodies but may modulate simultaneous binding of ACE2 and allosteric interactions between the binding sites.

To compare the free energy changes induced by mutations of the binding epitope we specifically examined the alanine mutation scanning profiles residues in different complexes (Supporting Information, Figure S3). By mapping positions of the binding epitopes onto these distributions, we observed high destabilization changes for RBM sites involved in interactions with B38 (Supporting Information, Figure S3A). Noticeably, binding free energy loss upon alanine mutations of K417 and N501 sites were quite appreciable, confirming that K417 is one of the central epitope residues for B38. A narrower spectrum of the binding epitope residues is evident in the complex with class II P2B-2F6 antibody (Supporting Information, Figure S3B).

Alanine scanning of only several epitope residues (Y449, L452, E484) produced large destabilization changes. The distribution also highlighted that the RBM residues near Y449 and E484 contribute appreciably to binding affinity by forming two small hotspot clusters (Supporting Information, Figure S3B). Alanine modifications in K417 and N501 produced only small changes with this class of antibodies. Consistent with the experimental deep mutagenesis, we observed a wide spectrum of large destabilization changes caused by alanine substitutions of the binding epitope residues in the complex with class III EY6A antibody (Supporting Information, Figure S3C). A particularly strong hotspot cluster is formed near conserved rigid positions F377, C379, Y380 and V382. The large destabilization changes in these sites are largely determined by the loss of protein stability upon alanine modifications of these hydrophobic residues. It should be noted that alanine mutations of K417, E484 and N501 yielded only very minor changes as these sites retain a considerable degree of mobility in the complexes with this class of antibodies (Supporting Information, Figure S3C). In general, these distributions highlighted the diversity of the binding energy hotspots, particularly indicating relatively moderate contribution of functional centers subjected to circulating mutational variants that displayed a fair amount of structural and energetic plasticity. Only for class II antibody P2B-2F6 E484 emerged as an important binding energy hotspot, while for other classes of antibodies single mutations in these functional positions typically incurred only relatively moderate destabilization changes. Nonetheless, a compounded effect of circulating mutations K417N and N501Y could be more significant for binding with class I B38 antibody. We argue that escape mutations constrained by the requirements for host cell binding and RBD stability may select energetically adaptable sites that compromise antibody recognition through modulation of global motions and allosteric protein responses.

### Perturbation-Based Scanning of Spike Residues in the SARS-CoV-2 S-RBD Complexes with Antibodies : Allosteric Duality of Functional Centers Targeted by Global Variants

Using the PRS method we probed the allosteric effector and sensor potential of the S-RBD residues in complexes with studied antibodies. In the perturbation-based scanning model, the effector profiles evaluate allosteric propensity of protein residues to efficiently propagate signals over long-range in response to systematically applied external perturbations. Accordingly, the local maxima along the effector profile may serve as an indicator of allosteric hotspots that can influence dynamic changes in other residues and may control signal transmission in the system. The sensor profile and respective distribution peaks highlight residues that have a strong propensity to sense signals and produce allosteric response through altered dynamics. We compared the PRS effector profile for the unbound S-RBD form and antibody-bound forms (Figure 9). In the complex with B38, several effector peaks corresponding to structurally stable RBD regions (residues 348-352, 400-406) are preserved in the bound form (Figure 9A). Due to B38-induced modulation of the effector profile, K417 and N501 positions become aligned with the notable peaks, suggesting that these residues acquire a strong allosteric potential in the complex and correspond to the stable effector hotspots (Figure 9A). On the other hand, the effector potential value for the E484 site remained largely unchanged as compared to the unbound form. The PRS sensor profile for the RBD-B38 complex (Figure 10A) indicated that none of the functional sites of circulating mutations are mapped onto distribution peaks. As a result, allosteric communication in this complex may be highly dependent mostly on two functional positions K417 and N501.

**Figure 9.**
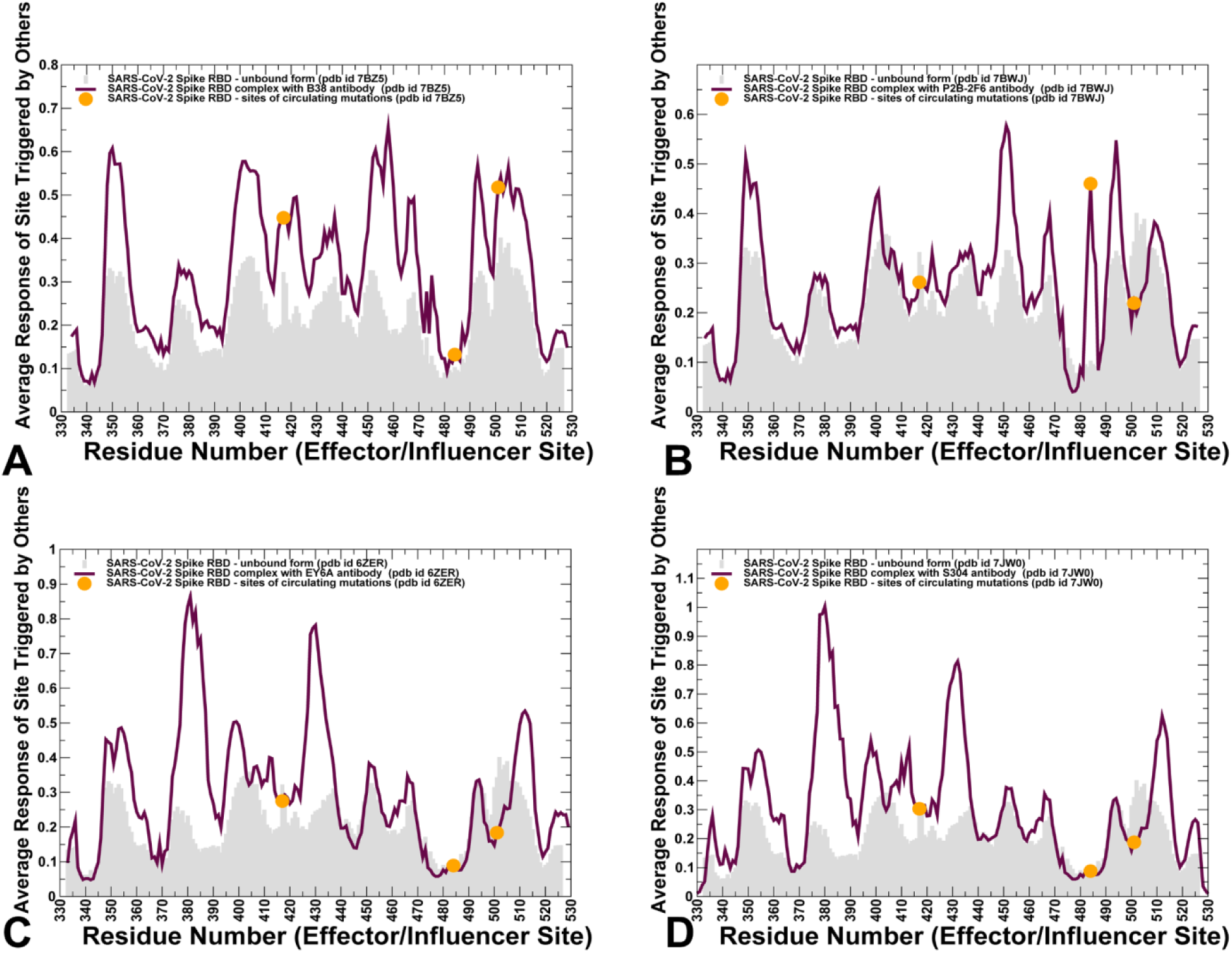
The PRS effector profiles for the SARS-CoV-2 S-RBD complexes with different classes of antibodies. (A) The PRS effector profile for the SARS-CoV-2 S-RBD complex with class I B38 antibody (pdb id 7BZ5). (B) The PRS effector profile for the SARS-CoV-2 S-RBD complex with P2B-2F6 antibody (pdb id 7BWJ). (C,D) The PRS effector profile for the SARS-CoV-2 S-RBD complexes with class III EY6A and S304 antibodies. The PRS effector profile for the unbound form of the S-RBD is shown on panels (A-D) in light grey bars. The PRS effector profiles for the SARS-CoV-2 S-RBD complexes are shown in maroon-colored lines. The position of functional RBD sites K417, E484, and N501 targeted by global circulating variants are highlighted in filled orange circles.

**Figure 10.**
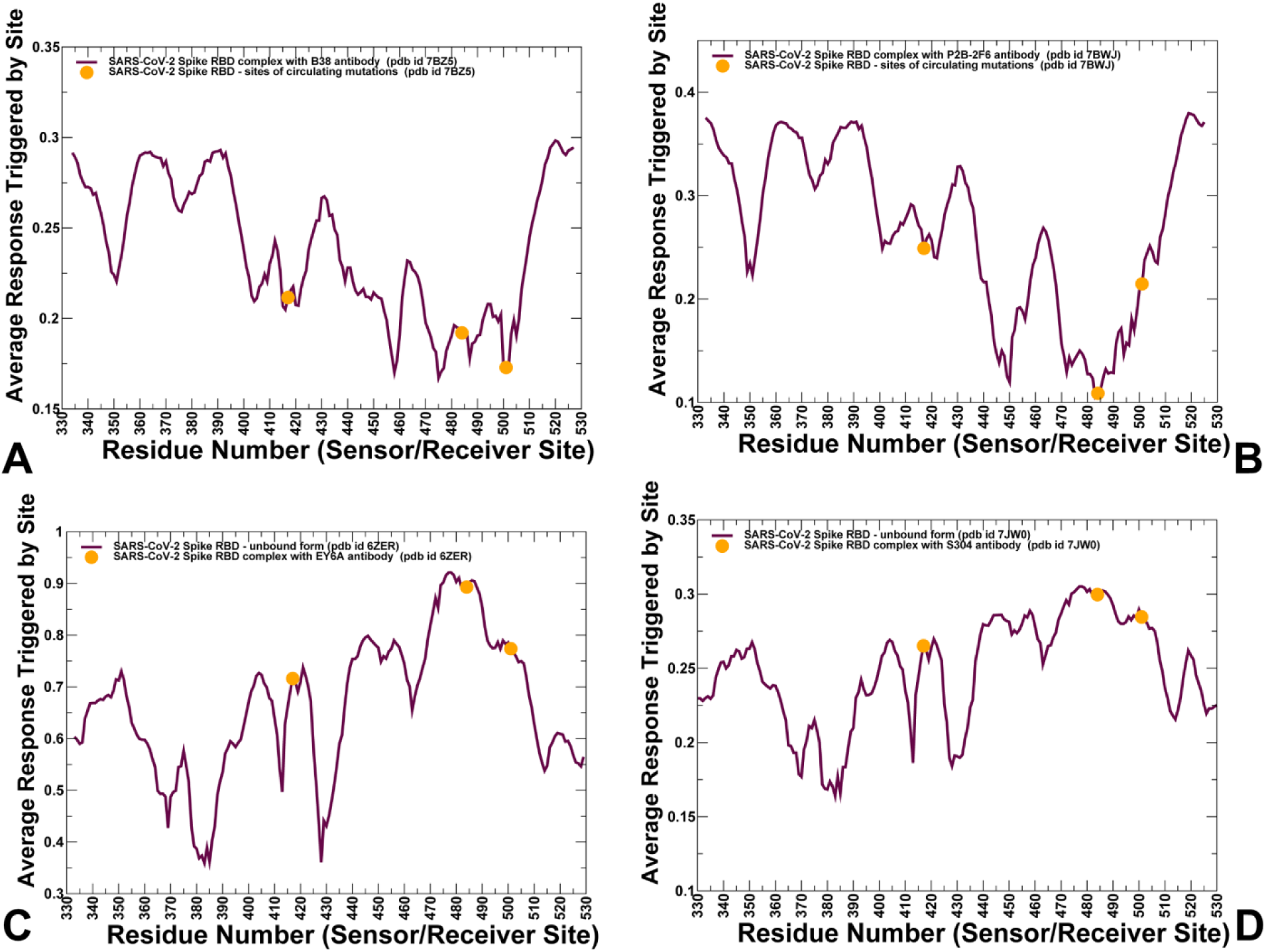
The PRS sensor profiles for the SARS-CoV-2 S-RBD complexes with different classes of antibodies. (A) The PRS sensor profile for the SARS-CoV-2 S-RBD complex with class I B38 antibody (pdb id 7BZ5). (B) The PRS sensor profile for the SARS-CoV-2 S-RBD complex with P2B-2F6 antibody (pdb id 7BWJ). (C,D) The PRS sensor profile for the SARS-CoV-2 S-RBD complexes with class III EY6A and S304 antibodies. The PRS sensor profile for the unbound form of the S-RBD is shown on panels (A-D) in light grey bars. The PRS sensor profiles for the SARS-CoV-2 S-RBD complexes are shown in maroon-colored lines. The position of functional RBD sites K417, E484, and N501 targeted by global circulating variants are highlighted in filled orange circles.

These observations complemented collective dynamics analysis showing that K417 and N501 residues become aligned with the hinge sites and allosteric effector centers that coordinate long-range communication between S-RBD and B38 molecules. While modifications of K417 and N501 residues appeared to trigger moderate changes in the binding affinity, the perturbations inflicted on these sites would have a significant effect on dynamics in other regions and allosteric signaling in the complex. In other words, allosteric hotspots may be energetically adaptable and not necessarily correspond to the structurally stable binding energy hotspots. These results support the recent studies suggesting that functional plasticity is central to allosteric regulation where allosteric hotspots may often correspond to structurally adaptable and moderately conserved protein positions.^100^

In the S-RBD complex with the class II antibody P2B-2F6, the effector profile was similar to the unbound form, but featured notable changes near E484 and N501 positions (Figure 9B). The allosteric effector propensities were markedly increased for E484 residue revealing a strong peak of the distribution, while the effector potential for N501 residue was clearly reduced (Figure 9B). Hence, antibody-induced stabilization of Y449 and E484 sites could lead to modulation of its allosteric propensities. As a result, these sites are not only important binding energy hotspots but also acquire significant allosteric potential in the complex. Although the sensor profile for this complex is similar to the one obtained for RBD-B38 complex, we noticed the increased sensor potential for K417 and N501 positions (Figure 10B). This suggested that functional sites may be engaged in an allosteric cross-talk of effector and sensor centers, where the binding signal induced by the P2B-2F6 antibody can be transmitted to other RBD regions.

In general, our findings suggested that class II antibody binding exemplified by P2B-2F6 may exhibit strong dependence on Y449 and E484 sites as mutations in these positions could significantly compromise both binding affinity and antibody-induced allosteric signaling. This may be an example where antibody binding recruits structurally adaptable E484 site and significantly alters the pattern of collective motions and communications in the spike protein. These results are consistent with the experimental evidence according to which E484 and F486 play a central role in this epitope.^32^

A different pattern of the effector and sensor centers emerged from the analysis of the S-RBD binding with EY6A and S304 antibodies. For both these III antibodies, the effector profiles were very similar and generally followed the distribution shape of the unbound S-RBD form (Figure 9C,D). We noticed a markedly increased effector peaks in the structurally stable regions of the RBD core that are further rigidified by EY6A and S304 binding. We capitalized on the deep mutational screening of CR3022 antibody^45^ that features exactly the same footprint as EY6A and S304. The sites of maximum escape for this class III antibodies correspond to C361, V362, K378, V382, S383, F392, T430, I434, A435 residues that are aligned with the effector peaks of the PRS distribution (Figure 9C,D). Accordingly, these sites could feature a significant allosteric potential and correspond also to the allosteric control centers in the complexes. Notably, none of the functional sites of circulating mutations were among effector centers. Moreover, the effector potential for these residues was relatively low, owing in part to the flexible nature of these regions that are solvent-exposed in complexes with EY6A and S304 (Figure 9C,D). The PRS sensor profiles for EY6A and S304 bound complexes showed strong peaks for K417, E484, and N501 residues (Figure 10C,D). Of notice is the dominant peak in the region centered on E484 and F486 residues. It implies that these positions correspond to the dominant sensor centers that can effectively “absorb” signals sent from the RBD core and execute protein response by altering collective dynamics in flexible regions.

In the PRS approach, the effector and sensor centers may act cooperatively to elicit long-range protein response to antibody binding and propagate allosteric signal through the protein. Together with the conformational dynamics analysis, our results also showed that class III antibodies may induce rigidification of the RBD core that is accompanied by the increased flexibility of the RBM region, which is a manifestation of dynamics-driven allosteric changes. This may explain the experimental results according to which even though EY6A cannot prevent ACE2 binding it can substantially increase the dissociation rate between ACE2 and RBD.^36^ This may reflect the increased mobility of the RBM residues and localization of key sensor hotspots in the regions interacting with the host receptor. Hence, functional sites K417, E484 and N501 can be involved as either effector or sensor hotspots in the complexes with different classes of antibodies. Hence, structurally and functionally adaptable allosteric centers may be targeted by mutational variants because even small perturbations in these positions would propagate over long-range allowing for modulation of antibody-induced protein response. Our findings showed that E484 site may be a critical effector hotspot for binding of class II antibodies, while serving as a dominant sensor center in response to binding of class III antibodies. Allosteric duality of this functional site could make it vulnerable to mutations which may alter collective dynamics and potentially be a driver of antibody resistance to the antibody classes I and II. Indeed, mutations in the epitope centered around E484 position (F486, F490) were shown to strongly affect neutralization for different classes of antibodies.^49, 50^ Collectively, our findings suggested that SARS-CoV-2 S protein may exploit plasticity of specific allosteric hotspots to generate escape mutants that alter response to antibody binding without compromising activity of the spike protein.

## Conclusions

We combined MD simulations with mutational and perturbation-based scanning approaches to perform a comprehensive structural profiling of binding and allosteric propensities of the SARS-CoV-2 S-RBD residues in complexes with three different classes of antibodies. Our analysis indicated that different classes of antibodies can uniquely change the spectrum of slow modes and induce specific functional motions. The EY6A and S304 antibodies targeting the cryptic binding site can modulate conformational dynamics without dramatically altering allosteric protein response but rather fine-tune dynamic changes at the ACE2-binding site. Consistent with deep mutational scanning experiments, in silico mutational profiling highlighted the diversity of the binding energy hotspots induced by different classes of antibodies. Mutational analysis of K417, E484 and N501 positions implicated in circulating variants showed that these residues correspond to the interacting centers with a significant degree of structural and energetic plasticity. Only for class II antibody P2B-2F6 one of these sites E484 is a prominent binding energy hotspot, while for other classes of antibodies single mutations in these functional positions typically incurred relatively moderate destabilization changes. Using perturbation-based scanning approach we probed allosteric propensity of spike residues and determined the allosteric hotspots involved in regulation of signaling. The results showed that sites of circulating mutations K417, E484 and N501 correspond to energetically adaptable allosteric hotspots that regulate functional motions and allosteric protein response to different classes of antibodies. This study provides a useful atomistic insight into allosteric regulatory mechanisms of SARS-CoV-2 S proteins, suggesting that circulating mutational variants are likely to emerge in structurally and evolutionary adaptable regulatory switch positions to elicit a differential protein response to the host cell receptor while evading antibody binding.

## AUTHOR INFORMATION

\***Corresponding Author** Phone: 714-516-4586; Fax: 714-532-6048;

The authors declare no competing financial interest.

## Acknowledgment

This work was partly supported by institutional funding from Chapman University. The author acknowledges support by the Kay Family Foundation Grant A20-0032.

## ABBREVIATIONS

SARS: Severe Acute Respiratory Syndrome
RBD: Receptor Binding Domain
ACE2: Angiotensin-Converting Enzyme 2 (ACE2)
NTD: N-terminal domain
RBD: receptor-binding domain
CTD1: C-terminal domain 1
CTD2: C-terminal domain 2

