## Supporting Figures S1-S3 for "Dynamic Profiling of Binding and Allosteric Propensities of the SARS-CoV-2 Spike Protein with Different Classes of Antibodies: Mutational and Perturbation-Based Scanning Reveal Allosteric Duality of Functionally Adaptable Hotspots"

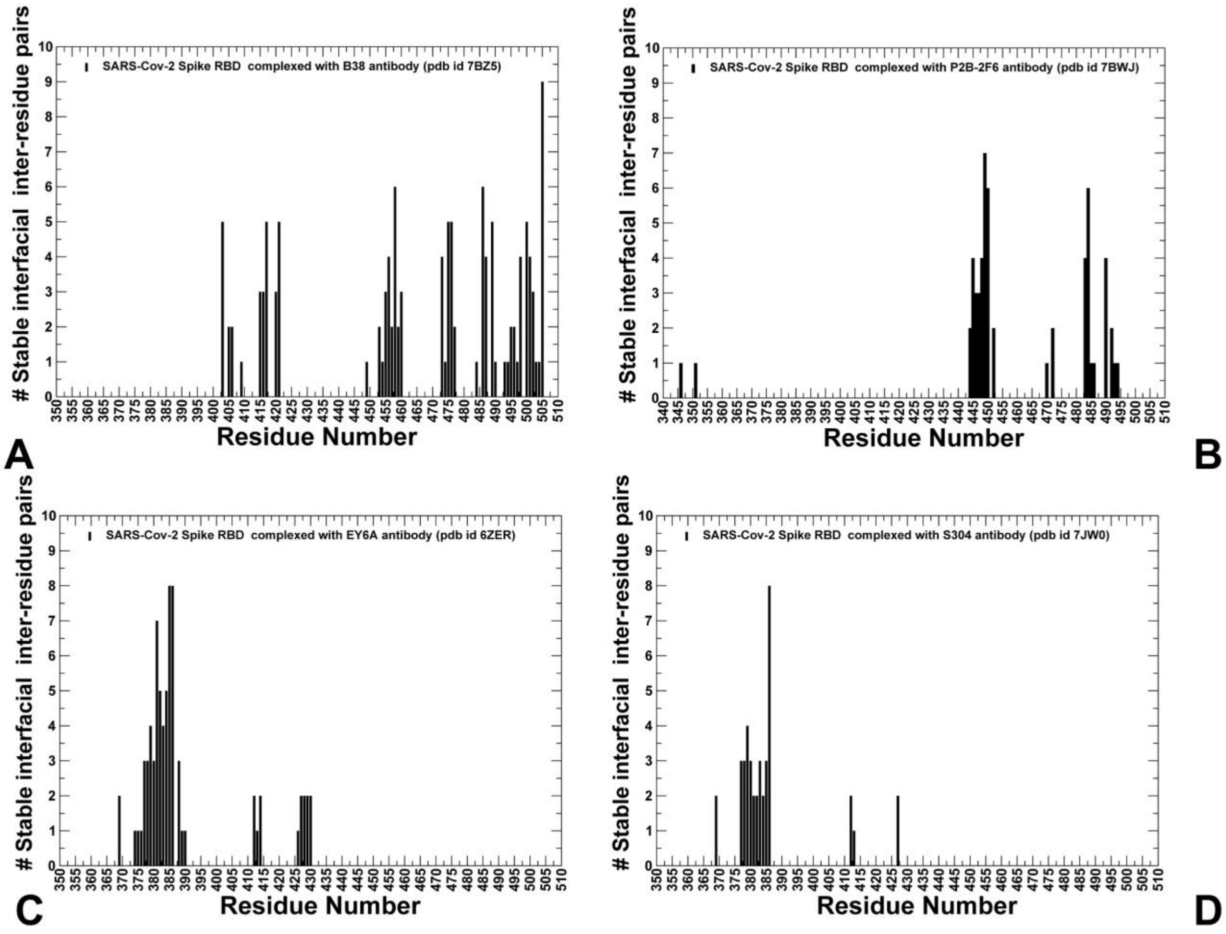


**Figure S1.** The distribution of the distinct inter-molecular residue pairs in the S-RBD complexes with class I B38 antibody (A), class II P2B-2F6 antibody (B), and class III EY6A (C) and S304 antibodies (D). The profiles are shown in black bars.

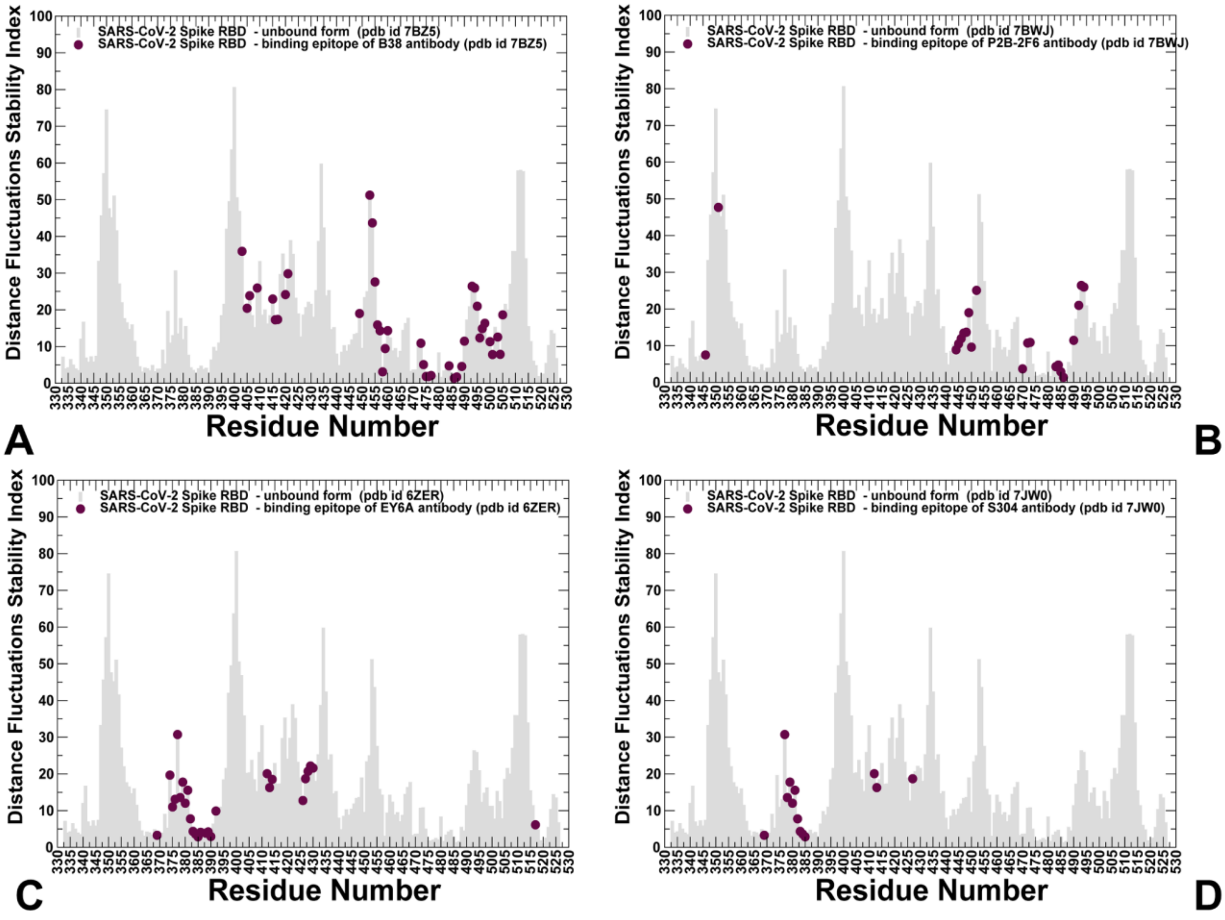


**Figure S2.** The distance fluctuation stability analysis of the SARS-CoV-2 S-RBD in the unbound form. The ensemble-based distribution of the distance fluctuation stability index for the unbound SARS-CoV-2 S-RBD is shown in light grey bars. The binding epitope residues of the S-RBD complexes with class I B38 antibody (A), class II P2B-2F6 antibody (B), and class III EY6A (C) and S304 antibodies (D) are highlighted in maroon-colored filled circles.

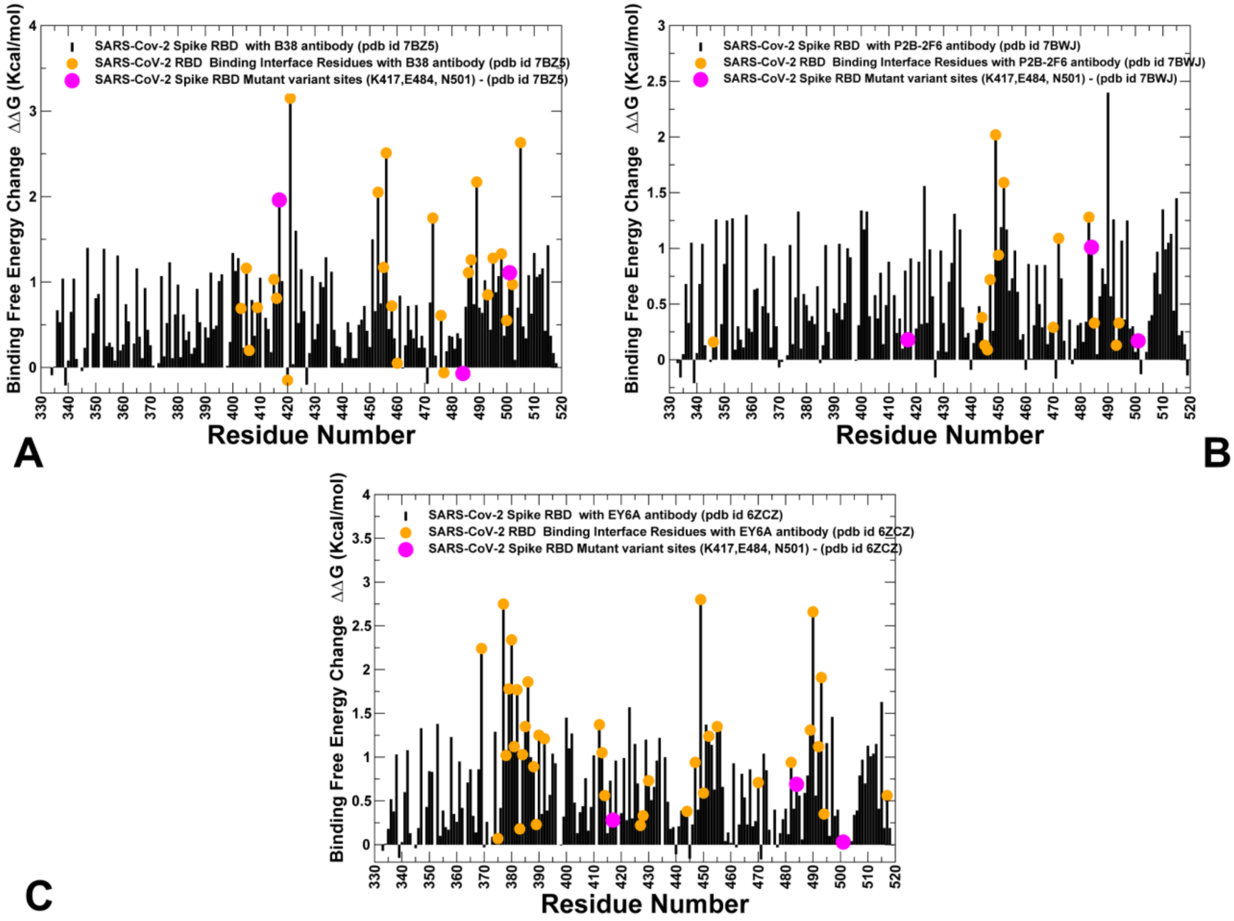


**Figure S3.** Alanine scanning of the RBD residues in the SARS-CoV-2 S-RBD in complexes with different classes of antibodies. (A) The binding free energy changes upon alanine mutations for the RBD residues in the SARS-CoV-2 S-RBD complex with class I B38 antibody (pdb id 7BZ5). (B) The binding free energy changes upon alanine mutations for the SARS-CoV-2 S-RBD residues in the complex with class II P2B-2F6 antibody (pdb id 7BWJ). (C) The binding free energy changes upon alanine mutations for the SARS-CoV-2 S-RBD residues in the complex with class III EY6A antibody (pdb id 6ZCZ). The binding energy changes for the protein residues are shown in maroon bars. The binding interface residues are depicted in orange filled circles and functional residues K417, E484 and N501 targeted by mutational variants are highlighted in magenta filled circles.
